# The representational nature of spatio-temporal recurrent processing in visual object recognition

**DOI:** 10.1101/2024.07.30.605751

**Authors:** Siying Xie, Johannes Singer, Bati Yilmaz, Daniel Kaiser, Radoslaw M. Cichy

## Abstract

The human brain orchestrates object vision through an interplay of feedforward processing in concert with recurrent processing. However, where, when and how recurrent processing contributes to visual processing is incompletely understood due to the difficulties in teasing apart feedforward and recurrent processing. We combined a backward masking paradigm with multivariate analysis on EEG and fMRI data to isolate and characterize the nature of recurrent processing. We find that recurrent processing substantially shapes visual representations across the ventral visual stream, starting early on at around 100ms in early visual cortex (EVC) and in two later phases of around 175 and 300ms in lateral occipital cortex (LOC), adding persistent rather than transient neural dynamics to visual processing. Using deep neural network models for comparison with the brain, we show that recurrence changes the feature format in LOC from predominantly mid-level to more high-level features. Finally, we show that recurrence is mediated by four distinct spectro-temporal neural components in EVC and LOC, which span the theta to beta frequency range. Together, our results reveal the nature and mechanisms of the effects of recurrent processing on the visual representations in the human brain.

## Introduction

Human visual object recognition is orchestrated by the interplay of feedforward and recurrent computations. Anatomically, this is evident by the number of feedforward and feedback connections complementing each other in visual circuits ^1–3^. Functionally, the feedforward sweep brings in information from the retina, enabling core object recognition through basic visual analysis ^4,5^. Then, the recurrent computations begin right after the first influx of feedforward information into the cortex. Recurrent activity contributes to object recognition not only when the viewing conditions are challenging ^6–12^, but also when objects are in plain view ^13–15^.

While the existence and importance of both feedforward and recurrent computations in object recognition is undoubted, their exact nature, i.e., where, when and how they affect visual processing remains incompletely understood ^16–19^. This is partly because their empirical dissection is challenging: shortly after the first feedforward sweep, feedforward and recurrent activity overlap in space and time ^20–22^, hindering their unique characterization.

Here, we used the classical experimental protocol of backward masking ^23–26^ to isolate the role of recurrent from feedforward activity ^27–31^. In backward masking a salient visual mask is shown shortly after a target image, impacting recurrent activity related to the target while leaving feedforward activity unaffected ^28,32–34^. Thus, the comparison of brain activity when participants view masked versus unmasked target images isolates the contribution of recurrent activity.

We recorded human brain activity with EEG and fMRI to resolve visual responses in time and space when a set of naturalistic object stimuli were either backward masked or not. We then used multivariate pattern analysis ^35–37^ to recover the neural representations of the image contents under the different masking regimes across time and space.

Comparing the neural activity related to the target images in the masked and unmasked conditions, we determined where, when and how recurrent activity contributes to human object vision. We first identified and characterized the spatio-temporal dynamics of visual recurrent activity. We then determined its respective spectral bases using time-frequency decomposition ^38–41^, and finally resolved its resulting visual feature format by relating neural representations to artificial neural network ^42–44^.

## Results

We presented 24 images of everyday objects on real-world backgrounds (Fig. 1A) to human participants while recording their brain activity with EEG (N = 31) and fMRI (N = 27) in separate sessions. On each trial, the target image was backward masked in one of two masking conditions: early mask or late mask (Fig. 1B). In the early mask condition, a dynamic mask rapidly followed the target after 17ms. The rapid succession of target and mask yields effective backward masking that disrupts recurrent processing ^28,32–34^. In contrast, in the late mask condition, the mask appeared after a delay of 600ms, leaving recurrent processing unaffected across an extended time window while otherwise keeping the stimulation across the whole trial the same.

**Figure 1.**
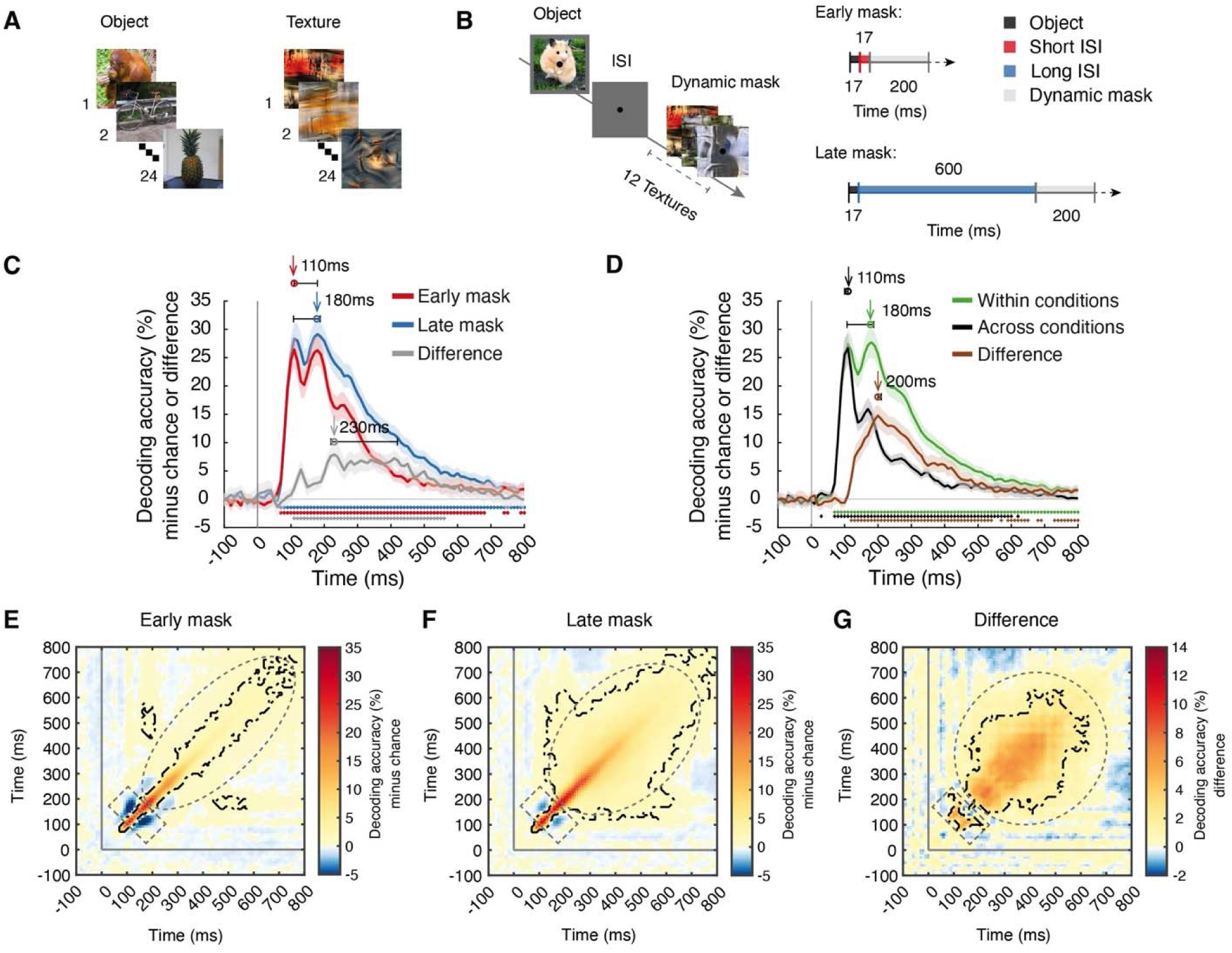
Experimental design and temporal dynamics of visual object representations. **(A)** Stimulus set. We used 24 real-world object images on natural backgrounds as target stimuli and 24 synthesized image textures created from an additional set of real-world object images for dynamic masks. **(B)** Experimental paradigm and timing parameters. On each trial, a briefly shown target object image was backward masked by a dynamic mask (i.e., a sequence of image textures) in one of two conditions: the early mask condition (short 17ms ISI) and the late mask condition (long 600ms ISI). **(C)** Results of object identity decoding in the early mask (red) and late mask (blue) conditions and their difference (gray). **(D)** Results of object identity decoding within (green) and across (black) masking conditions and their difference (brown). For **(C, D)**, chance level is 50%; significant above-chance level decoding is denoted by colored asterisks at the corresponding time points (N = 31, p < 0.05, right-tailed permutation tests, cluster definition threshold p < 0.005, cluster-threshold p < 0.05, 10,000 permutations); vertical gray line at 0ms indicates stimulus onset; shaded margins of time courses indicate 95% confidence intervals of the decoding performance determined by bootstrapping (1,000 iterations); horizontal error bars indicate 95% confidence intervals for peak latencies. **(E-G**) Results of time-generalized decoding object identity in the **(E)** early mask condition, **(F)** late mask condition, and **(G)** the difference. For **(E-G)**, chance level is 50%; time-point combinations with significantly above-chance level decoding are outlined in black dashed lines (N = 31, right-tailed permutation tests, cluster definition threshold p < 0.005, cluster-threshold p < 0.05, 10,000 permutations); vertical and horizontal gray lines indicate stimulus onset.

We used a multivariate pattern analysis framework to assess visual object representations captured by EEG and fMRI ^36,45^ to classify the objects in the target images from brain data. Because target images and masks were statistically independent by design across trials, classifying objects isolated the neural activity that related to the target from neural activity related to the mask.

We then characterized and compared object representations across the early mask and late mask conditions, revealing the temporal, spatial, and spectral characteristics as well as the representational format of the recurrent aspects of visual processing.

### The temporal dynamics of recurrent visual activity

To reveal the temporal dynamics of object representations in the early mask and late mask conditions, we conducted time-resolved multivariate pattern classification of object identity using EEG data. Classifying between all pairs of the 24 object images and averaging across pairs yielded a grand average object decoding time course for both masking conditions (Fig. 1C, for statistical details, see Supplementary Table 1). We assessed statistical significance using cluster-based inference (N = 31, right-tailed permutation tests, cluster definition threshold p < 0.005, cluster-threshold p < 0.05, 10,000 permutations), and report peak latencies as time points at which objects are best discriminated by neural representations with 95% confidence intervals derived by bootstrapping (1,000 samples) in brackets.

We observed a qualitatively similar and typical results pattern ^45,46^ in both masking conditions. Decoding accuracies fluctuated around baseline until 70ms after image onset, when they steeply rose to two peaks at ∼100ms and ∼200ms. The peak latencies for the objects in the early mask condition (110ms [110 – 180ms]) and the late mask condition (180ms [110 – 190ms]) coincided with the first and second peak respectively, without being significantly different (p > 0.05, Supplementary Table 1). This result demonstrates the presence of robust visual information in both masking conditions, warranting further analysis.

Comparing the decoding performance between the two masking conditions, we observed higher decoding in the late mask condition emerging after the first decoding peak (Fig. 1C, gray curve, cluster 110 – 560ms, peak latency 230ms [220 – 420ms]). This pattern was also present when decoding objects across the categorical boundary defined by naturalness or animacy (Supplementary Fig. 1A, B, Supplementary Table 2). Together, this provides a first characterization of the timing of recurrent activity.

The modest difference in the time-resolved decoding result patterns between the early and the late mask conditions might be interpreted as indicating a relatively minor role of recurrent processing in visual object processing. However, this conclusion is premature: similar overall time courses might hide qualitatively different visual representations across the two masking conditions.

To investigate whether the representations are strongly affected by recurrent processing, we decoded object identity across the two masking conditions ^47,48^. The rationale is that if visual representations are only weakly affected by recurrent processing, decoding results should be similar for the decoding within- and across-masking conditions. However, if recurrent processing affects visual representations more strongly, the across-condition decoding accuracy should be lower than the within condition accuracy. We found that cross-decoding was strongly reduced after the first peak (110ms [100 – 110ms], Fig. 1D, black curve) when compared to decoding within each masking condition (Fig. 1D, green curve, corresponds to the average of the blue and red curve in Fig. 1C). The difference between within- and across-conditions was significant after the first within-condition decoding peak (Fig. 1D, brown curve, clusters between 120ms and 800ms), with a peak at 200ms (200 – 210ms) (for statistical details, see Supplementary Table 1). This result pattern was also obtained when comparing within- and across-conditions decoding for training the classifiers on either the early or the late mask condition (Supplementary Fig. 1C, D, Supplementary Table 2). This indicates that recurrent processing strongly affects visual object representations after the first feedforward sweep from 120ms onward, thus detailing the temporal dynamics of recurrent processing.

If recurrent processing strongly affects visual object representations, the dynamics with which those representations emerge should also differ depending on the amount of recurrent activity involved. To assess this, we used temporal generalization analysis (TGA) ^49^ by decoding object identity across all time-point combinations in the EEG epoch. This resulted in time-time matrices for each masking condition (Fig. 1E, F) and their difference (Fig. 1G).

We observed similarities and differences for the two masking conditions. Concerning the similarities, in both masking conditions, significant effects were present from ∼70ms onwards, and decoding accuracies were highest close to the diagonal (i.e., similar time-points for training and testing), indicating that fast-evolving, transient representations dominate the neural dynamics. Further, we also observed significant off-diagonal generalization from 150ms on in both masking conditions, indicating the additional presence of stable and persistent representations. This shows that in both masking conditions, visual processing depends on both transient and persistent representations.

However, we also observed two key differences. For one, there was more widespread temporal generalization in the late mask than in the early mask condition (Fig. 1E, F, indicated by the length of the minor axis of the striped ellipses), and this difference was significant (Fig. 1G, striped ellipse). This suggests a stronger presence of persistent representations due to recurrent processing in the late mask condition. Second, we observed that below-chance decoding accuracies at the time-point combinations, i.e., ∼100ms and ∼200ms (Fig. 1E-G, striped rectangle), were lower in the early mask condition than in the late mask condition, emerging as a positive difference in their comparison (Fig. 1G, striped rectangle).

Negative decoding accuracies in off-diagonal regions of the TGA can be interpreted as an amplitude reversal of a time-locked (i.e., to the onset of the image) oscillatory neural component with a half-cycle length corresponding to the time difference between the time-points of the combination at which the negative decoding occurs ^50,51^ (for graphical illustration, see Supplementary Fig. 2). Assuming that in the early mask condition recurrent processing is reduced while feedforward processing is unaffected. This links feedforward activity to time-locked oscillatory components that are covered by time-varying recurrent activity in the late mask condition. In turn, in the early mask condition recurrent activity is reduced, and the time-locked feedforward-related oscillatory activity is uncovered. This result pattern was confirmed when comparing the decoding between within-condition of the late mask and the cross-decoding (Supplementary Fig. 3A-C), and it was reversed when comparing the decoding between the within-condition of the early mask and the cross-decoding (Supplementary Fig. 3D-F), supporting our interpretation.

Together, our results provide three key insights into the temporal dynamics of recurrent visual processing: firstly, recurrent processing affects visual object representations from ∼100ms onward, after the first feed-forward sweep, and most strongly around 200ms; secondly, it contributes specifically to the emergence of persistent representations; thirdly, it is less phase-locked to the onset of the stimulus than feed-forward activity.

### The spatial profile of recurrent visual activity

Next, we determined the spatial profile of recurrent processing across the visual brain. For this, we used an equivalent multivariate pattern analysis scheme and comparison strategy between masking conditions as for the temporal dynamics but applied in a spatially resolved way to fMRI data.

We focused on two regions of interest (ROI) in the visual ventral stream: the early visual cortex (i.e., V1, V2, and V3 combined) as the entry point of visual information in the cortex ^52,53^ and the lateral occipital complex (LOC) (Fig. 2A) as a central high-level hub for object representations ^54–56^. We decoded object identity in both masking conditions (Fig. 2B) as well as across masking conditions (Fig. 2C) and compared the results (N = 27, sign-permutation tests, FDR-corrected, p < 0.05).

**Figure 2.**
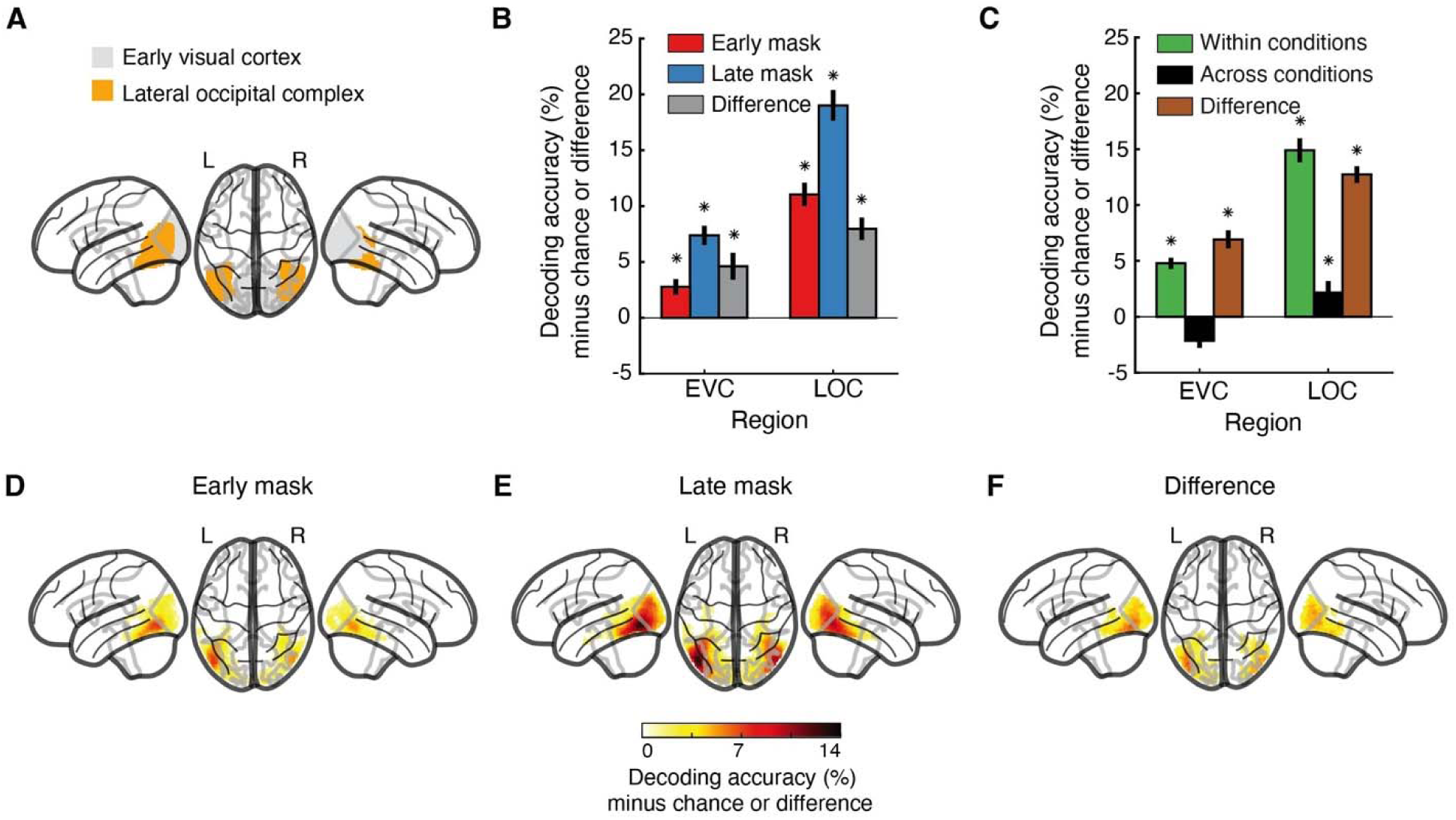
Cortical locus of visual object representations. **(A)** Visualization of the early visual cortex (EVC, i.e., V1, V2, and V3 combined) and the lateral occipital complex (LOC) ROIs. **(B)** Results of object identity decoding in the early mask condition, the late mask condition, and their difference. **(C)** Results of object identity decoding within and across masking conditions and their difference. For **(B, C)**, chance level is 50%; significant above-chance level decoding is denoted by black asterisks above the bars (N = 27, p < 0.05, right-tailed permutation tests, FDR-corrected); error bars indicate standard errors of the mean. **(D-F)** Results of the spatially unbiased searchlight decoding in the **(D)** early mask condition, **(E)** late mask condition, and **(F)** the difference. For **(D-F)**, chance level is 50%; only voxels with significant above-chance level decoding are shown (N = 27, right-tailed permutation tests, cluster definition threshold p < 0.005, cluster-threshold p < 0.05).

In line with the EEG results, there was above-chance decoding of object identity in both ROIs in both masking conditions (Fig. 2B, blue and red bars, all ROI-results FDR-corrected). Further comparing masking conditions, we found higher decoding accuracies for the late mask condition in EVC and LOC (Fig. 2B, gray bars), indicating that recurrent processing affects representations in both regions.

Akin to the EEG analysis, we next determined the degree to which recurrent activity alters visual representations. For this, we compared the within-condition decoding results (Fig. 2C, black bars) to the across-conditions results (Fig. 1D, green bars), noting their difference (Fig. 2C, brown bars). In both ROIs, the decoding accuracy was strongly reduced when decoding across masking conditions. In LOC, but not EVC, there was low but significant cross-decoding accuracy. An equivalent results pattern emerged when comparing within- and across-conditions decoding for training the classifiers on either the early or the late mask condition (Supplementary Fig. 4A, B). This indicates that recurrent activity strongly impacts visual representations in both EVC and LOC.

To explore the differences between the two masking conditions across the whole brain, we used a spatially unbiased fMRI searchlight analysis ^57,58^. Consistent with the ROI results, we found object identity information across the ventral visual stream in both masking conditions (Fig. 2D, E, right-tailed permutation tests, cluster definition threshold p < 0.005, cluster-threshold p < 0.05, 5,000 permutations). Comparing decoding in the early mask versus the late mask conditions revealed widespread effects in the ventral stream with a maximum in the high-level ventral cortex (Fig. 2F). This reinforces the view that recurrent activity strongly affects visual representations across the ventral stream.

### Recurrent processing affects the format of visual representations

We next investigated how recurrent processing affects the format of visual representations. For this, we used representational similarity analysis (RSA) ^37,61^ to compare representations in the brain and in the layers of an 8-layer AlexNet deep neural network (DNN) model trained on object categorization ^59,60^ (Fig. 3A). The rationale is that correspondence to layers along the DNN hierarchy reveals the complexity of the representational format in the brain, from low-complexity features in the early layers to high-complexity features in the late layers ^42–44^.

**Figure 3.**
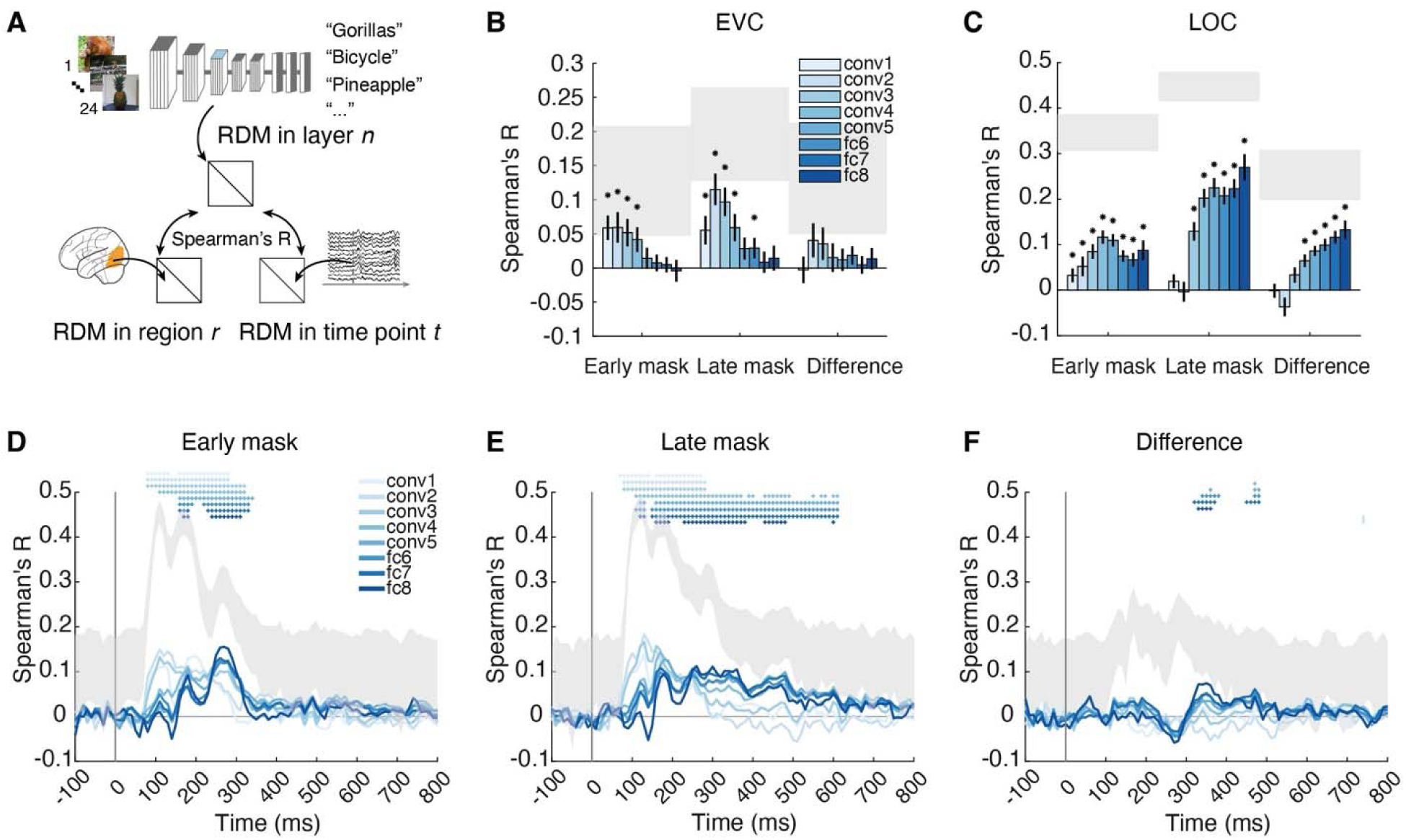
The representational format of visual representations resolved in space or time. **(A)** RSA linking brain responses to layer-wise activation patterns in a DNN model (AlexNet trained on object categorization). We obtained RDMs for each layer of the DNN, each ROI in fMRI, and each time point in EEG. We then compared (Spearman’s R) the DNN RDMs with the EEG and fMRI RDMs, respectively. **(B, C)** RSA results linking **(B)** EVC and **(C)** LOC to DNN layers. For **(B, C)**, significant correlations are marked by black asterisks above bars (N = 27, p<0.05, right-tailed permutation tests, FDR corrected); error bars depict standard errors of the mean; shaded gray areas indicate the noise ceiling. **(D-F)** RSA results linking DNNs to EEG in the **(D)** early mask condition, **(E)** late mask condition, and **(F)** difference therein. For **(D-F)**, significant correlations at time points are denoted by asterisks colored by layer (N = 31, right-tailed permutation tests, cluster definition threshold p < 0.005, cluster-threshold p < 0.05, 10,000 permutations); horizontal error bars indicate 95% confidence intervals for peak latencies, shaded gray areas represented the noise ceiling.

We began the investigation of the format of visual representations as localized in EVC and LOC using fMRI. In EVC, we identified the strongest correspondences with the early to middle DNN layers in both masking conditions (Fig. 3B). The differences between masking conditions were numerically most pronounced in the early layers, albeit not significantly. This suggests that feedforward and recurrent processing in EVC primarily involve the processing of low-level features. A supplementary analysis that compared the visual representations as revealed by the within- and across-conditions decoding to the DNN model showed an equivalent result pattern (Supplementary Fig. 5A), further strengthening this view.

In contrast, in LOC, we made two key observations. First, while there were correspondences with the middle to deep layers in both masking conditions (Fig. 3C), there was a shift in the peak correspondence from the highest layer (i.e., layer 8) in the late mask condition to a middle layer (i.e., layer 4) in the early mask condition. Correspondingly, the comparison of results between masking conditions revealed differences in mid-to-late layers, with a peak in the latest layer. Second, there was correspondence in LOC to early layers (i.e., layers 1-2) in the early mask condition but not in the late mask condition. Both observations were also present when comparing within- and across-conditions decoding results (Supplementary Fig. 5B). This suggests that in LOC the representational format shifts from lower to higher complexity through recurrent activity.

Next, we assessed the change in the representational format of visual representations across time using EEG. We observed correspondence to all layers of the DNN in both masking conditions (Fig. 3D, E) with a temporal progression in peak correspondence from lower layers early in time to higher layers later in time ^62,63^ (for statistical details, see Supplementary Table 3). This shows that in both masking conditions, visual representations emerge along a cascaded processing hierarchy characterized by increasing feature complexity ^5,17,64,65^. To assess the feature complexity and timing of recurrent processing directly, we determined the difference in correspondence between the masking conditions (Fig. 3F). We found that the difference was highest in the middle and high layers between ∼300ms and 500ms. This indicates that recurrent activity changes the representational format to one of higher complexity features. Consistent with this conclusion, equivalent results patterns were observed in a supplementary analysis comparing the visual representations revealed by the within- to across-conditions decoding to the DNN model (Supplementary Fig. 5D).

Finally, for both EEG- and fMRI-based analyses, we confirmed the main results pattern using another DNN architecture (i.e., ResNet50 ^66^, Supplementary Fig. 6), indicating the generalizability of the conclusions across models.

Together, this shows that recurrent processing leaves the format of EVC representations unaffected in terms of visual feature complexity. In contrast, recurrent processing changes the format of LOC and late representations from lower to higher complexity, revealing the nature of its effect on the representational format.

### The spatiotemporal dynamics of changes in representational format through recurrence

Visual processing evolves dynamically across spatial locations in the brain and across time simultaneously, necessitating a spatiotemporally resolved view ^35,67^. However, the analyses so far assessed visual representations and their format separately in space and time. For a fully spatio-temporally resolved view, we used RSA-based commonality analysis ^68,69^ (Fig. 4A), providing time courses of shared variance with each DNN layer in EVC and LOC (for statistical details, see Supplementary Table 5).

**Figure 4.**
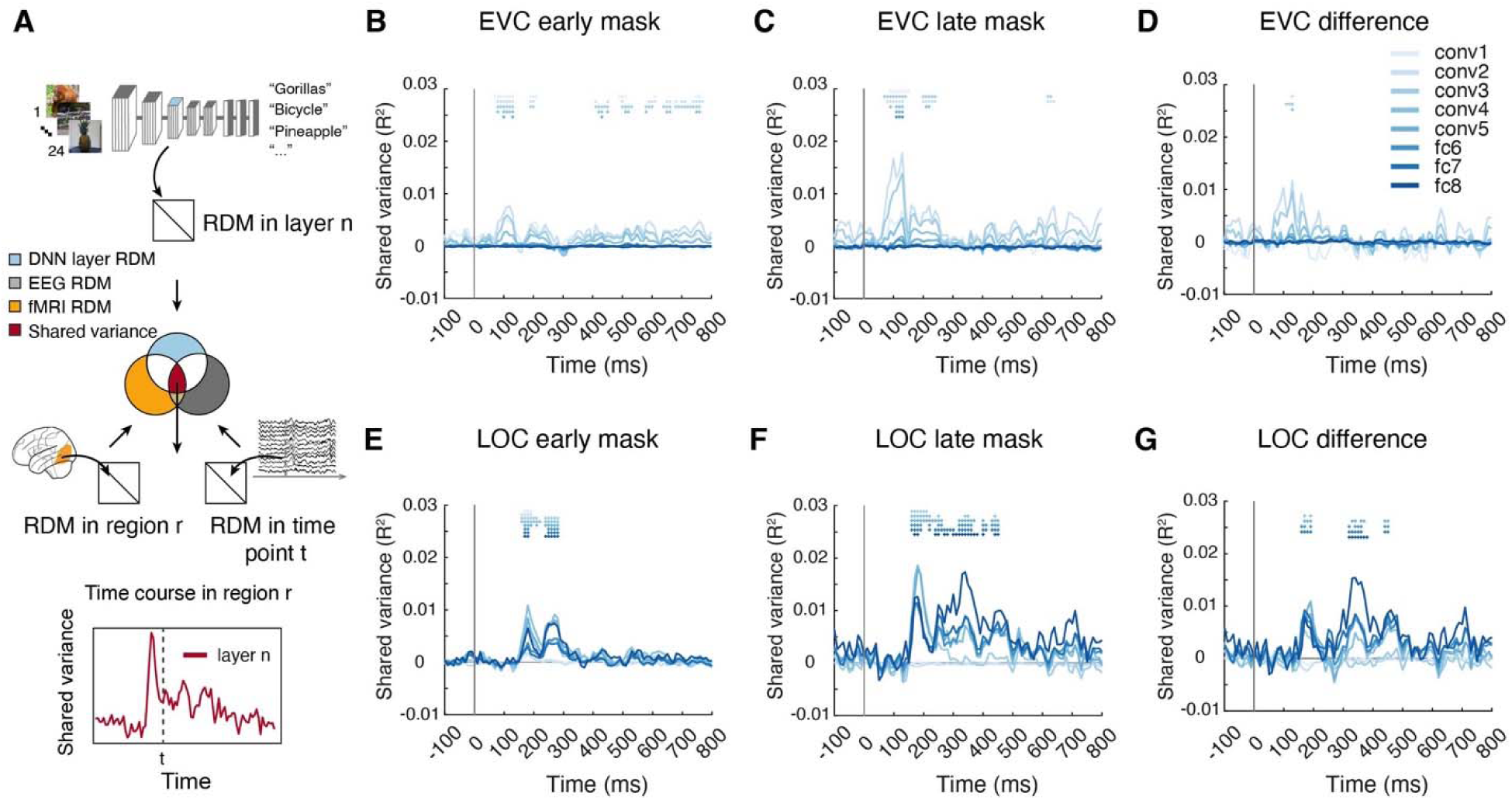
The format of spatiotemporally resolved visual representations. **(A)** Commonality analysis based on RSA, linking temporal dynamics (EEG), cortical locus (fMRI) and feature complexity (DNN layers of AlexNet). This yielded time courses of shared variance for each DNN layer in EVC and LOC respectively (here: layer 3 in LOC). **(B-G)** Time courses of shared variance with DNN features in the **(B, E)** early mask condition, **(C, F)** late mask condition, and **(D, G)** difference between them, in EVC **(B-D)** and LOC **(E-G)** respectively. For **(B-G)**, significant effects at time points are denoted by asterisks color-coded by DNN layer (N = 31, right-tailed permutation tests, cluster definition threshold p < 0.005, cluster-threshold p < 0.05, 10,000 permutations).

In EVC, we observed an emergence of visual representations of low- to mid-complexity with peaks early in time, predominantly at 120 – 130ms, both in the early mask condition and the late mask condition (DNN layers 1-6, Fig. 4B, C). The difference between masking conditions emerged early (peaks at ∼90 – 130ms) and was in low-to-middle complexity, too (DNN layers 1-5, Fig. 4D). This shows that recurrent activity impacts visual representations in EVC early in time and in a low-to-mid-complexity format.

In LOC, we observed the emergence of visual representations of all complexity levels at a later stage than in EVC, with two peaks at ∼ 200ms and 300ms in both masking conditions (Fig. 4E, F). The difference between masking conditions was in features of middle-to-high complexity (DNN layers 4-8, Fig. 4G). This shows that recurrent activity impacts visual representations in LOC later in time and in a mid-to-high complexity format.

In sum, recurrent activity modulates EVC representations early in processing in low-to-mid complexity format, and LOC representations later in processing in mid-to-high complexity format.

### The spectro-temporal basis of recurrent processing

The transmission of visual information and the formation of visual representations ^70,71^ is fundamentally indexed by oscillatory neural activity. Based on previous work in human and non-human primates, we hypothesized that recurrent processing should be evident in the low-frequency range between the theta- and the beta-range ^72–74^. Thus, in the next step, we investigated the spectral characteristics of visual processing in the early mask condition and the late mask condition. For this, we decoded object identity from EEG data resolved both in time and frequency (Fig. 5A), considering power and phase of the signals separately.

**Figure 5.**
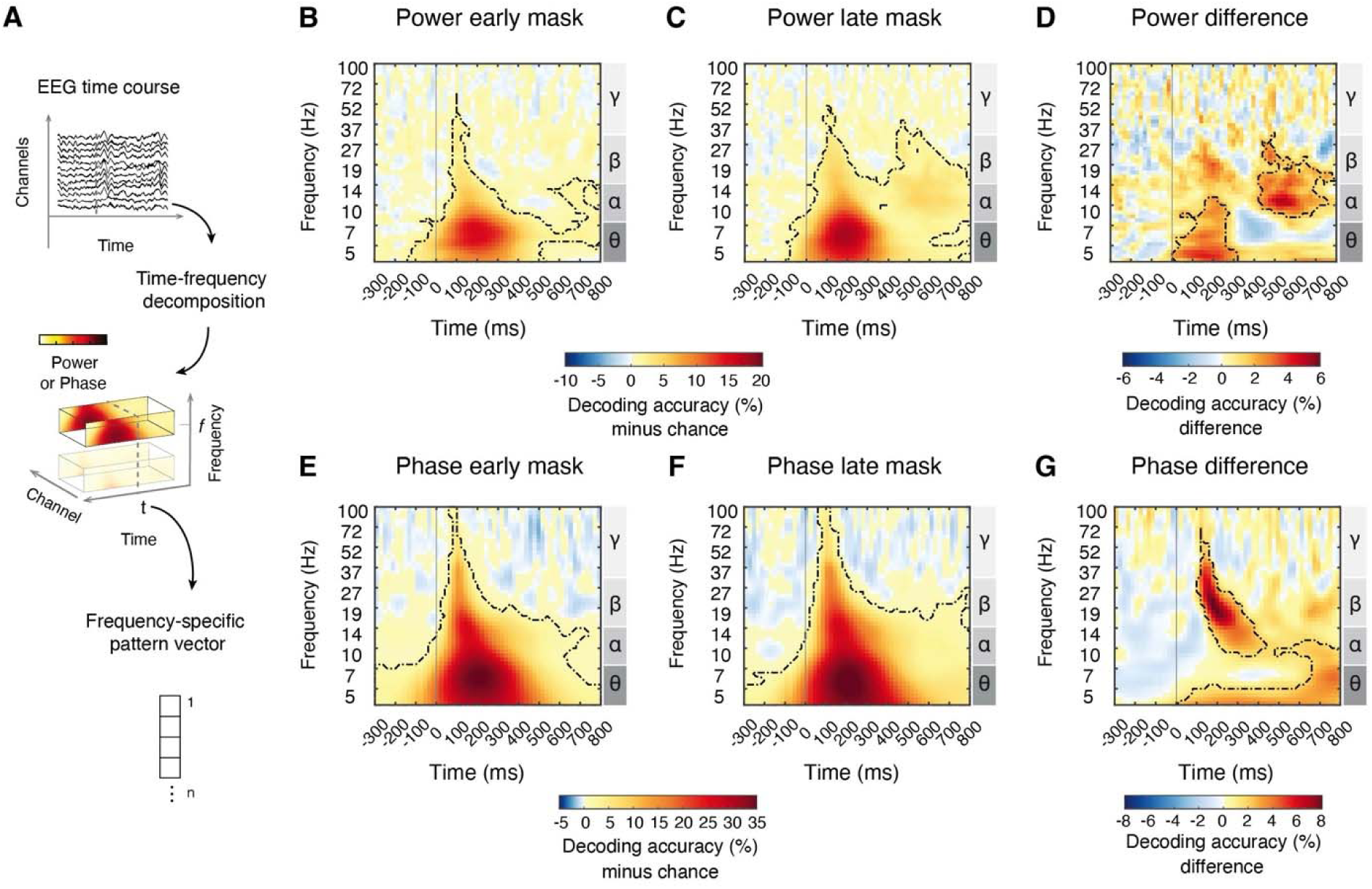
Spectral characteristics of visual representations. **(A)** Using time-frequency decomposition we extracted frequency-specific response pattern vectors across EEG channels for power (63) and phase values (63 × 2 = 126) separately. **(B-G)** Results of time- and frequency-resolved object identity decoding in the **(B, E)** early mask condition and **(C, F)** late mask condition, and **(D, G)** difference between them, based on power values **(B-D)** and phase values **(E-G)**. For (**B-G)**, chance level was 50%; time-frequency combinations with significant above-chance decoding are outlined by black dash lines (N = 31, right-tailed permutation tests, cluster definition p < 0.05, significance p < 0.05, 10,000 permutations); the vertical gray line indicates stimulus onset, and the right y-axis labels indicate frequency bands.

Across both masking conditions and for both power (Fig. 5B, C) and phase (Fig. 5E, F), we observed significant object decoding in a broad frequency range. The decoding peak was consistently within the theta band (∼6 Hz) at ∼200ms (for statistical details see Supplementary Table 6). This establishes the sensitivity of the analysis and warrants further inspection by contrasting the masking conditions.

Comparing the results of the early mask condition to the late mask condition, we observed four components with distinct temporal and spectral characteristics (Fig. 5D, G; for statistical details see Supplementary Table 6). Two clusters were in the power domain (Fig. 5D) and two in the phase domain (Fig. 5G). In detail, in the power domain, there was a cluster before 300ms in the theta-alpha frequency range (peak at 4.27 Hz, 160ms), and a later cluster after 400ms in the alpha-beta frequency range (peak at 10.72 Hz, ∼540ms). In the phase domain, there was a cluster between 100ms and 400ms in the alpha-beta frequency range (peak at 19.35 Hz, 200ms) and a cluster in the theta range across the entire temporal range after stimulus onset (peak at 10.03 Hz, 560ms). A supplementary analysis comparing the within- and across-conditions decoding (Supplementary Fig. 7) revealed a more widespread effect that largely encompassed the clusters observed here.

Together, this establishes the spectro-temporal basis underlying recurrent visual processing as four distinct components with specific spectro-temporal profiles.

### Feature format and cortical origin of the spectral components underlying recurrent processing

In a final step, we asked for each of the four spectro-temporally identified components: what is their spatial origin in the brain, and in what feature format do they represent object information? To address both questions, we again used RSA-based commonality analysis ^68,69^, relating frequency-based EEG signals to fMRI signals from EVC and LOC and layers of the DNN model (Fig. 6A).

**Figure 6.**
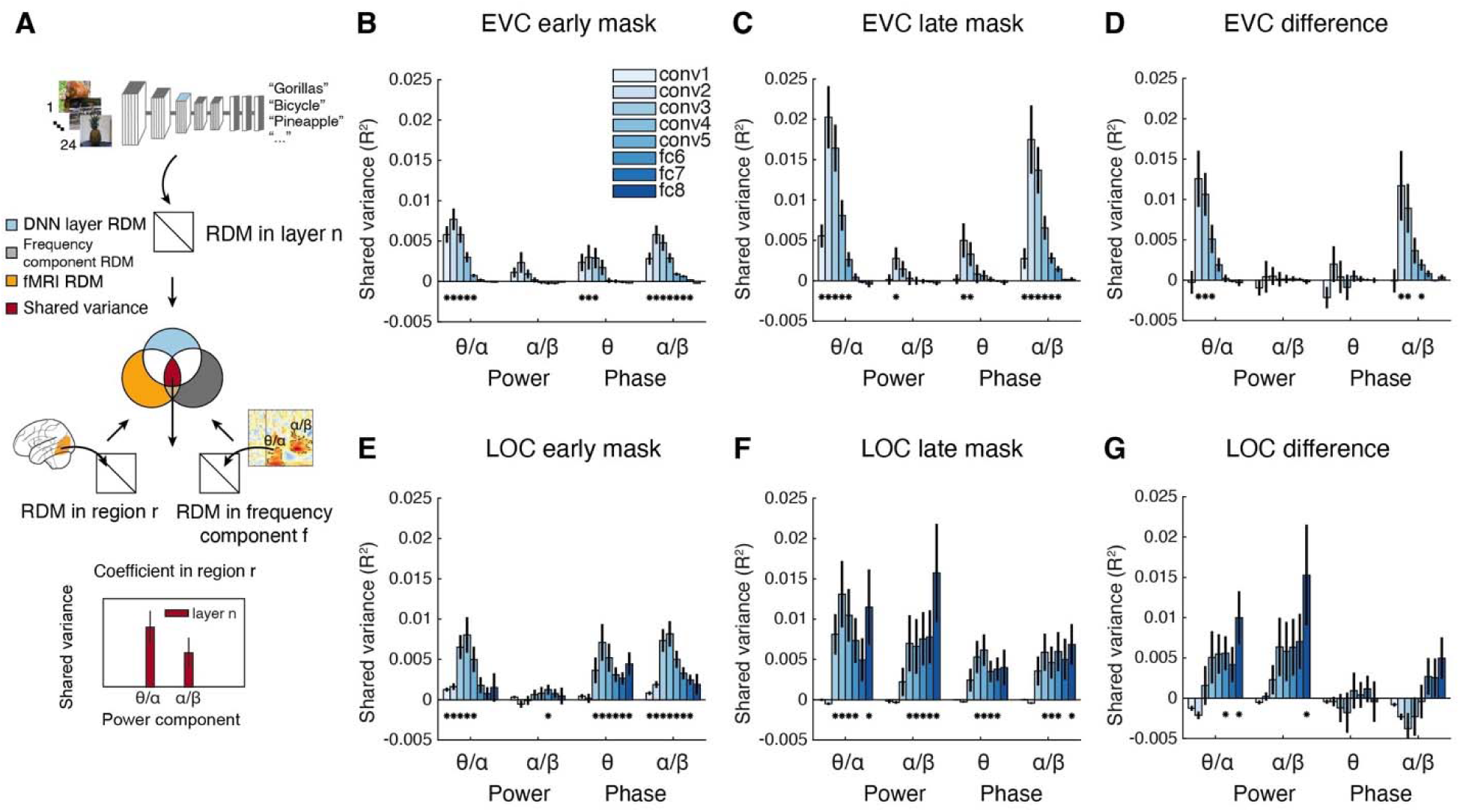
Spatial profile and feature format of the spectral components underlying recurrent processing. **(A)** Commonality analysis based on RSA linking identified time-frequency resolved dynamics (EEG), cortical locus (fMRI) and feature complexity (DNN layers of AlexNet). This analysis yielded coefficients of shared variance for each of the four identified components and for each DNN layer in EVC and LOC respectively. **(B-G)** Shared variance in the **(B, E)** early mask condition, **(C, F)** late mask condition, and **(D, G)** difference between them, in EVC **(B-D)** and LOC **(E-G)** respectively. For **(B-G)**, significant effects at DNN layers are denoted by asterisks (N = 31, right-tailed permutation tests, cluster definition threshold p < 0.005, cluster-threshold p < 0.05, 10,000 permutations).

We first note results common to all four components, forming the basis for further discriminative investigation. First, we observed significant relationships to DNN layers for all components, regions, and both masking conditions (Fig. 6 B, C and Fig. 6 E, F; except for EVC and the alpha-beta power component in the early mask condition, see Fig. 6B), demonstrating the analytical feasibility of the approach. Second, similar to the previous analyses, the shared variance was generally lower in the early mask condition compared to the late mask condition.

However, key differences emerged when isolating recurrent processing by considering the difference between the early and late mask conditions with respect to the feature complexity of visual representations (Fig. 6D, G). Concerning EVC (Fig. 6D), we found a relationship between the two components emerging early in time, that is, the theta-alpha power component and the alpha-beta phase component, but not for the other two clusters. This relationship was strongest in low-and mid-level visual features. In contrast, concerning LOC (Fig. 6G), we found a relationship between the two power-defined components, one early in time and the other late in time, strongest for high-level visual features.

Together, these results reveal the spectro-temporal basis of recurrent processing in EVC and LOC by comprehensively characterizing its distinct spectrally identified components, in terms of their specific feature complexity and temporal profile.

## Discussion

### Summary

We combined a backward masking paradigm with multivariate analysis on EEG and fMRI data, along with computational modelling, to characterize when, where and how recurrent processing affects object representations. Harvesting the detailed structure of visual representations beyond grand-average responses to visual stimulation, we showed that recurrence substantially affects the image-specific geometry of visual representations.

First, regarding the spatiotemporal dynamics, we found that recurrence affects visual representations across the ventral visual stream, early on at ∼100ms in EVC and in two later phases of ∼175 and 300ms in LOC, adding persistent rather than transient neural dynamics to visual processing. Next, we determined the feature complexity and spectral basis of the effect of recurrence on visual representations. We found that recurrence changes the feature format in LOC from mid- to high-level features and is mediated by four distinct spectro-temporal components in EVC and LOC in the theta to beta frequency range.

### The spatio-temporal dynamics of recurrent processing

Our separate analyses of EEG and fMRI data revealed a broad impact of recurrent processing: it affects visual representations starting at 100–120ms, with a peak at 200ms in a wide plateau, and across the ventral visual stream.

The combination of EEG and fMRI dissected these broad effects into distinct components for EVC and LOC. In EVC, recurrence affected visual representations early with a peak at 100ms. This is in the range of previously observed early effects of recurrence in non-human primate EVC ^75^, associated with contextual modulation and figure-ground segregation ^76–78^ that originate from within the ventral visual stream rather than with attentional effects ^16,79,80^.

In LOC, recurrence affected visual representations later, with two peaks at ∼175 and 300ms. The earlier peak at 175ms is consistent with effects of masking observed in humans invasively in V4/pIT ^32^, potentially originating from prefrontal cortex ^81,82^ and modulating visual activity in monkey V4 and IT ^83,84^. The later peak at 300ms might reflect pattern completion, as indicated by delayed responses in invasive studies of human IT in a similar time frame ^11,85^. The origin of this late effect might be medial temporal lobe regions such as parahippocampal cortex that activates as early as 270ms after stimulus onset ^86–88^. Alternatively, attentional effects might be driving the late effect, consistent with reports of human and non-human attentional modulation in high-level ventral visual cortex starting at 150ms ^75,89–92^.

Our results cannot ultimately determine whether non-visual regions contributed to the observed effects as here, fMRI coverage was restricted to the ventral visual stream. Future research assessing the whole brain, including frontal ^84,93–95^ and parietal ^96–99^ regions, is needed.

Temporal generalization analysis added two further insights into the temporal dynamics of recurrent processing. For one, recurrence specifically contributed to the emergence of persistent, rather than transient representations. This is consistent with the observation that masking reduces firing duration in single cells in monkey IT ^100,101^, and that masking reduces persistence in the visual representations of occluded objects in humans ^9^. Together this supports the view that recurrence plays an active role in accruing and maintaining important information online for further processing and decision making ^45,102–106^.

The second insight is that recurrent activity is less phase-locked to the onset of the stimulus than feedforward activity. Notably, the spectral basis of recurrent activity in LOC is in power only (Fig. 6G), in contrast to EVC where it is in power and phase (Fig. 6D). This suggests an increase of variability in phase over the course of processing, possibly due to accruing variability as information propagates increasingly back and forth along the visual processing hierarchy.

### Recurrence transforms the feature format in LOC from mid- to high-level complexity

Using deep neural networks to model the representations from EEG and fMRI data ^42–44^, we found that recurrence changes the feature complexity of representations in LOC, but not in EVC.

In LOC, we observed a shift of representational format from predominantly mid-level to more high-level features through recurrent processing. This has three implications. First, it adds algorithmic specificity to the observations from invasive recordings in non-human primates that feature coding in high-level ventral visual cortex is dynamic, changing the code over time from global to fine-grained ^107^, individual object parts to multipart configuration ^108^, and from a code supporting detection to one for discrimination ^109^. Second, it qualifies the finding that masking affects firing rate and stimulus specificity in monkey IT ^31,100^, linking those observations to the lack of recurrent activity mediating high-complexity features ^7,84^. Finally, it converges with visual imagery and working memory studies indicating that recurrent processing carries high-complexity features ^41,110^. However, a limitation of our finding is that we cannot distinguish whether the observed effect indicates the addition of new features to LOC representations through recurrence that are absent in feedforward processing ^13^, or the modulation of the gain of already present features, e.g., through attention ^111–114^.

In contrast to LOC, we did not find evidence for a change in feature complexity in EVC from its low-level complexity format (Fig. 4D and Fig. 6D). Analogous to the case of LOC, this suggests two different mechanisms underlying recurrence in EVC. One is that recurrent activity in EVC amplifies features encoded already in the feed-forward sweep ^16^. The other is that it adds new features of low-level complexity, consistent with observations of dynamical feature coding in orientation and color ^115,116^ and changes to receptive field structure ^117^. To distinguish these potential mechanisms of recurrence in both LOC and EVC future work is needed, for example, investigating the finer-grained encoding of single features rather than feature complexity ^118,119^ and modulating attentional state ^120–122^.

Note that here we used DNNs as a tool to characterize feature complexity rather than to directly model human visual processing. Future work is needed that carefully and explicitly models how feedforward and recurrent activity ^18,123,124^ account for core object recognition ^13^, as well as visual behavior ^125^.

### The spectral basis of recurrent processing

Our results on the spectral basis of recurrent processing go beyond previous work in several ways: by identifying distinct oscillatory components of the spectral basis of recurrent processing, by linking those components differentially and directly to stimulus properties and by clarifying their distinct relationship to EVC and LOC as well as their distinct feature format ^39,126–128^.

We find that a set of distinct spectro-temporal components of power and phase in the theta to beta frequency range subserve recurrent processing. Our findings refine the view that low-frequency rhythms may generally serve as a neural index for recurrent processing ^73,74^ by showing that recurrent processes can further be subdivided into early recurrent processes (in the phase domain) that refine the representations of basic visual features, followed by later recurrent processes (in the power domain) that sculpt the representations of complex visual features in higher levels of the visual hierarchy (for a detailed discussion of each component, see Supplementary Discussion 1).

Our results further support the broad notion that theta ^129^, alpha ^74,130^ and beta ^72–74,131^ frequencies mediate recurrent activity and play an active role in cognition ^132–135^ and vision in particular^41,136–138^, rather than in inhibition of irrelevant information ^130,139^ or cortical idling ^140,141^.

### Backward masking as a tool to dissect recurrent processing

A key assumption on which our interpretations rest is that the difference between early and late mask conditions in neural activity isolates recurrent processing to a relevant degree. While not undoubted ^142,143^, this assumption is supported by a large number of studies linking backward masking to recurrent rather than feedforward processing ^20,28,32,34,144^, impacting the communication between and to visual regions ^34,84,145^.

Our results invite future backward masking studies employing multivariate analysis to further confirm and dissect the sources of recurrent activity identified here. This might in particular involve causal interventions such as TMS ^146^ to determine the sources of recurrent activity across cortex, and layer-specific fMRI analysis ^147–149^ to distinguish recurrent from feedforward processing based on cortical layers ^1,2,150^.

### Conclusion

In sum, recurrent activity substantially affects the ventral visual stream, first in EVC and subsequently in LOC. Recurrent processing drives a shift in the feature format of LOC from mid- to high-level complexity and is linked to distinct spectro-temporal components in the theta to the beta frequency range. These findings characterize where, when, and how recurrence affects visual representations, furthering the understanding of how the recurrent information flow in the brain mediates visual object perception.

## Materials and Methods

### Participants in EEG and fMRI experiments

We conducted two independent experiments: an EEG and an fMRI experiment. Thirty-two participants took part in the EEG experiment, of whom one was excluded due to high-frequency noise in the recordings (N = 31, mean age 26.6 years, standard deviation 4.8 years, 20 female). Twenty-eight participants took part in the MRI experiment, of whom one was excluded due to failure of the stimulus presentation equipment (N = 27, mean age 27.7 years, standard deviation 4.6 years, 19 female). There was an overlap of four participants between the EEG and the fMRI participant sample. All participants had normal or corrected-to-normal vision. The study was conducted according to the Declaration of Helsinki and approved by the local ethics committee of the Freie Universität Berlin.

### Stimulus set

The stimulus set consisted of a set of target object images and a set of image textures used to create dynamic object masks.

The set of target object images consisted of 24 object images (Fig. 1A). Each image showed an object of a different object category and was cropped quadratically to the size of the centrally presented object. The 24 object images were a subset of a larger set of 118 images ^151^. The rationale for selecting the stimulus subset was as follows. Brain responses to natural images are typically highly correlated across the stages of the visual processing hierarchy. That is, two images that elicit similar responses at one stage tend to elicit similar responses at another stage, too. This makes assessing the role of different processing stages and the information they send in a forward or backward direction using multivariate analysis methods particularly difficult: due to the high correlations observed, experimental effects cannot be uniquely assigned to particular stages. To improve the chances of eliciting dissociable responses across the visual processing hierarchy in our experiment, we selected the stimulus set that yielded low correlations between the entry (early visual cortex, EVC) and the endpoint (inferior temporal cortex, IT) of the ventral visual pathway. For this, we used fMRI data in EVC and IT for the 118-image superset from a previous experiment ^151^. We assessed the similarity of representations in EVC and IT for the 118 images using representational similarity analysis (RSA) ^37,61^. To select 24 images that yielded uncorrelated responses, we used a genetic algorithm ^152^ for optimization. In detail, the optimization constraint was to minimize the absolute value of correlation between EVC and IT representational dissimilarity matrices (RDMs). The RDMs for the chosen 24-stimulus set yielded the desired low similarity between EVC and IT (R = 0.0018) on the preexisting fMRI data set. In comparison, this was lower than a random selection of 24 stimuli would have been (as assessed by 1,000 random draws, average R = 0.211, standard deviation = 0.101).

We created a set of image textures to be used for dynamic backward masks. For this, we chose a different subset of 24 object images randomly from the 118-image set and converted the images to textures that conserved the low- and mid-level image statistics of the images without portraying identifiable objects ^153^. We next created 24 dynamic masks that consisted of a sequence of 12 textures each, by randomly assigning 12 of the 24 texture images in random order to a dynamic mask.

### Experimental procedures

#### Main experiment & experimental design

We presented object images to participants in a backward masking paradigm (Fig. 1B). The general experimental design, stimulus presentation parameters, and trial structure were equivalent in both the EEG and the fMRI experiments. We describe the crucial elements common to EEG and fMRI first before detailing the modality-specific differences.

On each trial, a single object image (referred to as “target”) was briefly displayed for 17ms, followed by a 200ms dynamic mask. Object images and dynamic masks were randomly paired for each trial. We manipulated the target’s visibility by varying the inter-stimulus interval (ISI) between target and mask. This defined two conditions: in the early mask condition, the ISI was 17ms; in the late mask condition, the ISI was 600ms. During each trial, one of the 24 dynamic masks was presented. Stimuli were presented centrally on a gray background with a size of 5 x 5 degrees visual angle, overlaid with a bull’s-eye fixation symbol with a diameter of 0.1-degree visual angle ^154^. The texture images of dynamic mask were positioned and sized identically to the target object images. Participants were instructed to fixate on the fixation symbol throughout the experiment. We used Psychophysics Toolbox ^155^ for experimental presentation.

#### EEG experimental procedures

In the EEG experiment, participants completed a total of 2,544 main trials partitioned into 26 blocks of 3.5 minutes each. Throughout the experiment, each object image was presented a total of 53 times in both the early mask condition and the late mask condition.

We assessed the participants’ recognition performance with additional task trials that were interspersed every 4 to 6 (average: 5) main trials. The task was to identify the object image in the previous trial from a pair of images in a two-alternative forced choice (2-AFC) task. For this, two images were presented side by side for 500ms: one of the images presented was the image from the previous trial, and the other image was randomly chosen from the remaining 23 images. Participants indicated their response with a button press.

Participants were instructed to refrain from blinking throughout the experiment except during the additional interspersed task trials, when participants were asked to blink when they gave their responses. While the inter-trial interval (ITI) between main trials was between 900ms and 1,100ms, following the 2-AFC trial, the ITI was extended to 2,000ms to prevent motor artifacts from influencing the EEG recordings of the subsequent trial.

Participants had high task performance in both masking conditions, suggesting that they attended to the stimuli even under viewing challenging conditions (for details and statistics, see Supplementary Table 7). Further, as expected, the task performance was worse for the early mask condition than for the late mask condition trials. This confirms the efficacy of the backward masking procedure in reducing object visibility.

#### fMRI experimental procedures

In the fMRI experiment, participants performed a total of 12 runs, each lasting 6.5 minutes. In each run, each object image was presented twice in the early mask condition and the late mask condition, resulting in 96 main trials per run. The trial-onset synchrony (TOA) was 3,000ms.

Main trials were interspersed with null trials (34 per run), during which only the background but no stimulus was shown.

Participants were instructed to attend to the object images and respond with a button press if an object image was repeated in two consecutive trials (i.e., a one-back task on the target images). Object repetitions occurred ten times per run.

As in the EEG experiment, participants had overall high task performance, with worse performance for the early mask condition than for the late mask condition trials (for details and statistics, see Supplementary Table 7).

#### fMRI localizer experiment

To define the regions-of-interest (ROIs) early visual cortex (EVC) and object-selective lateral occipital cortex (LOC), we performed a separate fMRI localizer run. The localizer run was conducted prior to the fMRI main experiment runs. The stimulus set comprised 40 images of objects and scrambled objects each.

The localizer run used a fMRI block design. Each block lasted 15 s. During each block, 20 stimuli were centrally presented within an area of 5 x 5 degrees visual angle at a rate of 650ms on and 100ms off. There were 6 object and scrambled object blocks each. They were presented in counterbalanced order and randomly interspersed with 7 baseline blocks during which only the background was shown.

Participants were instructed to fixate on a centrally presented fixation symbol that was presented throughout the experiment, and to respond to one-back repetitions of images with a button press. Repetitions occurred a total of 9 times over the course of the localizer experiment.

### EEG data acquisition, preprocessing, and time-frequency decomposition

We recorded EEG data using an ActiCap 64 electrodes system and a Brainvision actiChamp amplifier. 64 electrodes were placed according to the 10-10 system, with an additional ground electrode and a reference electrode placed on the scalp. The signals were sampled at a rate of 1,000 Hz and online filtered between 0.03 and 100 Hz. All electrodes’ impedances were kept below 10 kΩ during the recording.

We preprocessed EEG data offline using the Brainstorm-3 toolbox ^156^. We removed noisy channels (average 2.2 channels per participant, standard deviation 1.8 channels) identified through visual inspection. We then filtered the data with a low-pass filter at 40 Hz. Eyeblinks and eye movement artifacts were detected using independent component analysis (ICA). We visually inspected the resulting components and removed those resembling the spatial properties of eyeblinks and eye movements (average 2.7 components per participant, standard deviation 0.9 components). We segmented the continuous data in epochs between -200ms and 800ms with respect to target image onset and baseline-corrected the segmented data by subtracting the mean of the 200ms interval before stimulus onset from the entire epoch. We finally applied multivariate noise normalization on the preprocessed data to improve the signal-to-noise ratio and reliability of the data ^157^. This formed the data for the temporally resolved decoding analyses. For time-frequency analysis, we preprocessed the data again the same way except for two differences: 1) we did not apply offline filtering, and 2) we segmented the continuous data into longer epochs (-600ms to 1,200ms) to enable better estimation of signals at lower frequencies.

#### Time-frequency decomposition of the EEG data

We performed time-frequency decomposition by applying complex *Morlet* wavelets. The wavelets, resembling complex sine waves modified by a Gaussian function, covered frequencies from 4 to 100 Hz in 50 logarithmically spaced increments. The Gaussian taper characteristics varied across this frequency range, with temporal full-width-half-maximum (FWHM) ranging from 20ms to 500ms as frequency decreased and spectral FWHM ranging from 1Hz to 31Hz as frequency increased.

We applied the complex *Morlet* wavelets for each channel and each trial of the EEG data at 2ms intervals (i.e., 500Hz). At each time point, this yielded 50 distinct frequency coefficients corresponding to the range of 4 to 100 Hz. At each time-frequency point, we computed two measures: the power and phase of the oscillation. To determine the absolute power values, we took the square root of the resulting time-frequency coefficients. To determine the phase values, we determined the real (sine) and imaginary (cosine) components from the time-frequency coefficients. This decomposition procedure yielded frequency-resolved EEG signals to be used for further time-frequency resolved decoding analyses. To decrease computation time and disk space usage, we downsampled the time points of frequency-resolved signals at 20ms intervals after time-frequency decomposition.

### fMRI data acquisition, preprocessing and univariate analysis

We acquired T2* and T1-weighted MRI data using a 3T Siemens Tim Trio scanner with a 32-channel head coil. We acquired T2*-weighted BOLD images using a gradient-echo EPI sequence. The acquisition parameters were as follows: TR = 2,000ms, TE = 30ms, FOV = 224 x 224 mm^2^, matrix size = 112 x 112, voxel size = 2 x 2 x 2 mm^3^, flip angle = 70°, with 30 slices and a 20% gap. The acquisition volume covered the occipital and temporal lobes and was oriented parallel to the inferior temporal cortex. Additionally, we obtained a T1-weighted image for each participant as an anatomical reference (MPRAGE; TR = 1,900ms, TE = 2.52ms, TI = 900ms, matrix size = 256 x 256, voxel size = 1 x 1 x 1 mm^3^, and 176 slices).

We performed fMRI data preprocessing using SPM12 (https://www.fil.ion.ucl.ac.uk/spm/software/spm12/). This involved realignment, slice-time correction, co-registration to the anatomical image, and normalization to MNI space. For the fMRI data of the localizer experiment, but not the main experiment, we additionally applied smoothing with a Gaussian kernel (FWHM = 5 mm). For the fMRI data from the main experiment, we additionally estimated noise components using the Tapas PhysIO toolbox ^158,159^ by creating tissue-probability maps from each participant’s anatomical image and extracting noise components from the white matter and CSF maps combined with the fMRI time series.

We used a general linear model (GLM) to estimate responses for the 48 experimental conditions (i.e., the 24 object images presented in either the early mask condition or the late mask condition). The analysis was conducted in a participant-specific fashion. We applied the GLM estimation to the preprocessed fMRI data for each run. We entered experimental condition onsets and durations as regressors into the GLM. Nuisance regressors comprised noise components and movement parameters. We evaluated 20 different GLMs by convolving regressors with 20 distinct hemodynamics response functions (HRFs) as derived from a large fMRI dataset ^160^. For each voxel, we then identified the HRF that resulted in the lowest average residual ^161^ and chose the corresponding estimates for further analysis. This approach resulted in 48 beta maps (one for each experimental condition) for each run and participant.

We used a separate GLM to estimate responses for the localizer run. We included block onsets and durations as regressors for the 3 conditions (i.e., objects, scrambled objects, and baseline), along with movement parameters as nuisance regressors. We convolved the regressors with the canonical HRF. We computed two contrasts from the resulting GLM parameter estimates that were used at a later step for voxel selection in the ROI analysis. The first contrast was defined as object + scrambled objects > baseline to define EVC. The second contrast was defined as objects > scrambled objects to define LOC. This yielded two t-value maps for the localizer run per participant.

#### Definition of fMRI regions of interest (ROIs)

For each participant, we identified two regions of interest (ROIs) within the ventral visual stream: early visual cortex (EVC) and lateral occipital complex (LOC). To determine the boundaries of these ROIs, we used participant-specific t-value maps from the localizer run threshold at p < 0.0001 intersected with anatomical masks. For the EVC definition, we intersected the thresholded t-value map (object + scrambled objects > baseline) with the combined anatomical region masks of V1, V2, and V3 obtained from the Glasser Brain Atlas ^162^. For the LOC definition, we intersected the thresholded t-value map (objects > scrambled objects) with a mask of LOC derived from a functional atlas ^163^. We removed any voxels shared between the EVC and LOC ROIs to avoid overlap. This process resulted in the definitions of two ROIs for each participant.

### Multivariate pattern analysis on EEG and fMRI data

An analytical challenge in comparing neural activity evoked by target images versus target image with a backward mask is the confounding effect introduced by the mask. Previous studies addressed this challenge by using subtraction design, for example, by including trials showing only the mask and subtracting the resulting neural activity from the neural activity evoked by the stimulus plus mask ^28,164^. Here, instead, we used a content-sensitive multivariate pattern analysis on EEG and fMRI data to dissect neural activity of the target image from neural activity evoked by the mask. The rationale is that in our design, target and mask stimuli were statistically independent, so multivariate pattern analysis classifying target object images revealed neural activity related to object images rather than the mask.

We performed multivariate pattern analysis on EEG and fMRI data using linear support vector machines ^165^ as implemented in the LIBSVM toolbox ^166^ in MATLAB (2021a). We conducted all analyses on a participant-specific basis.

#### Temporally resolved decoding analysis from EEG data

To determine when the brain processes object information, we conducted a time-resolved decoding analysis ^45,167^. We examined EEG data from -200ms to 800ms with respect to target image onset, in 10ms intervals. At each time point, we extracted trial-specific EEG channel activations and arranged them into 64-dimensional pattern vectors for each of the 24 object image conditions for each masking condition, separately. We conducted two types of analysis: within- and across-masking conditions object decoding.

In the within-masking condition analysis, we separately decoded object conditions for the early mask and the late mask conditions. For each of the 24 image conditions, we first randomly grouped trials into four equally sized bins and averaged them to create four pseudo-trials to enhance the signal-to-noise ratio (SNR). Employing a leave-one-out cross-validation approach, we then divided these pseudo-trials into training (three pseudo trials) and testing sets (one pseudo trial) to pairwise decode object identity. We then decoded object conditions pairwise for all object condition combinations. The resulting decoding accuracies were arranged into a 24 × 24 decoding accuracy matrix, with rows and columns corresponding to the decoded object conditions. This matrix is symmetric across the diagonal, with the diagonal being undefined. We repeated this analysis 100 times, randomly assigning trials to pseudo-trials each time. Averaging results over repetitions yielded one 24 × 24 decoding accuracy matrix for each time point, separately for the early and late mask conditions.

In the across-masking conditions analysis, we proceeded accordingly but assigned pseudo-trials to the training set and testing set from different masking conditions. That is, we trained on data recorded in the early mask condition and tested on data from the late mask condition (or vice versa). We averaged the results across both training and testing directions. This yielded one 24 × 24 decoding accuracy matrix for each time point.

In both analyses, averaging across the 24 × 24 entries of decoding accuracy at each time point resulted in a grand-average decoding accuracy time course.

#### Time generalization decoding analysis

We used time-generalization decoding analysis to determine how visual representations relate to each other across different time points. We proceeded as for the within masking condition time-resolved decoding analysis, except that classifiers trained on data from a particular time point were tested iteratively on data from all other time points. The rationale here is that successful generalization across time points indicates the similarity of visual representations over time. This analysis yielded 24 × 24 decoding accuracy matrices for each combination of time points from - 200 to +800ms. By averaging the entries of each decoding accuracy matrix across time point combinations, we obtained a temporal generalization matrix (TGM), where rows and columns are indexed by training and testing time points, respectively.

#### Time-frequency-resolved decoding analysis from EEG frequency power and phase

To determine the spectral properties of visual object representations in the two masking conditions, we conducted a time-frequency-resolved decoding analysis. This analysis was identical to the time-resolved analysis described above, but instead of decoding from raw activation values, we decoded object identity from patterns of power or phase value. We performed the analysis separately for 50 frequency bins spanning from 4 Hz to 100 Hz, using either power or phase values. In the power-based analysis, decoding was based on 64 power values corresponding to the 64 EEG channels. For the phase-based analysis, decoding used 128 values corresponding to the concatenation of the 64 sine and 64 cosine values. This resulted in one 24 × 24 decoding accuracy matrix for each time point and frequency bin, for the power- and phase-based analyses. Averaging across the 24 × 24 entries of decoding accuracy resulted in a grand average time-frequency matrix, where time points and frequency bins are indexed in rows and columns, respectively.

#### Spatially resolved decoding analysis from fMRI data

We conducted two types of decoding analyses on the fMRI data: region-of-interest (ROI)-based and spatially unbiased volumetric searchlight-based decoding ^57,58^ on the fMRI data.

For the ROI-based analysis, we arranged beta values from voxels of a given ROI into pattern vectors for each of the 24 experimental conditions and each of the 12 runs of the main fMRI experiment. To enhance signal-to-noise ratio, we grouped 3 runs into 4 bins and averaged across runs, creating four pseudo-run fMRI pattern vectors ^168^. Then for each ROI, we performed object decoding on these pseudo-run fMRI pattern vectors in a leave-one-pseudo-run-out manner. Averaging across iterations yielded a 24 × 24 decoding accuracy matrix for each ROI, participant, and masking condition.

For the searchlight-based analysis, for each voxel in the 3D fMRI volume, we defined spheres of voxels around it with a radius of four voxels. For each sphere, we arranged voxel values into pattern vectors. We then decoded object identity as described for the ROI-based analysis. This yielded a 24 × 24 decoding accuracy matrix for each voxel in the 3D fMRI volume for each participant, and each masking condition.

In both ROI and searchlight-based analyses, averaging across the 24 × 24 entries of decoding accuracy resulted in either a single value or a 3D map of grand average decoding accuracy, respectively.

### Representational similarity analysis (RSA)

RSA is a framework to relate representations across different measurement and signal spaces, such as those defined by different brain imaging modalities (EEG and fMRI) or computational models ^37,61^. The idea is to abstract from incommensurate measurement spaces into a common similarity space where representations can be directly compared.

For each masking condition, the analysis proceeded in two steps. In the first step, within each signal space of interest (e.g., fMRI responses in ROI, EEG broadband responses at particular time points, EEG spectral responses at time-frequency combinations, and activations of DNN layers), we calculated the dissimilarity between condition-specific multivariate activity patterns for all pairwise combinations of the 24 object conditions. We aggregated the results in representational dissimilarity matrices (RDMs), where rows and columns were indexed by the 24 object conditions. These RDMs summarize the representational geometry within each signal space. In the second step, we compared the RDMs across signal spaces using Spearman correlations, yielding a measure of their similarity. We provide the details for each of the two steps below.

#### Step 1: Construction of RDMs

For the brain data, we used the decoding accuracy matrices resulting from the decoding analyses detailed above as RDMs. This yielded RDMs a) from the temporally resolved EEG decoding analysis for each time point, b) from the time-frequency-resolved EEG decoding analysis for every time-point and frequency combination, separately for power and phase, and c) from the spatially resolved fMRI decoding analysis for each ROI.

For the computational model, we built RDMs from an AlexNet architecture trained for object categorization on the ImageNet dataset ^59,60^. AlexNet is an 8-layer deep neural network (DNN) commonly used as a baseline for brain-DNN comparisons ^169^. Using the MatConvNet toolbox ^170^, we fed our object stimuli into the pre-trained AlexNet and extracted the activation patterns for each stimulus from each of the five convolutional layers (conv1 to conv5) and the three fully connected layer (fc6, fc7, and fc8).

To test the generalizability of our conclusion across different DNN models, we also built RDMs using the ResNet50 architecture ^66^, pre-trained on the ImageNet dataset ^59^ for object categorization. ResNet50 features a distinct architecture compared to AlexNet, consisting of an initial convolutional layer followed by four residual blocks, each containing multiple convolutional layers with skip connections, and leading to a final classification layer. We fed the object stimuli into ResNet50 and extracted the activation patterns for each stimulus from the last layer of each of the four residual blocks (block1 to block4) as well as from the final classification layer (fc).

We quantified the dissimilarity of the activation patterns by calculating 1-Pearson’s R for each pair of stimuli. This resulted in eight RDMs for AlexNet layers and five RDMs for ResNet50 layers.

#### Step 2a: Standard RSA - relating DNN RDMs to EEG and fMRI RDMs

To characterize the format of neural representations, we related DNN RDMs from each layer to EEG and fMRI RDMs (Fig. 3a). The idea is that ascending layers of a DNN capture features of increasing complexity. Thus, relating neural representations to each DNN layer informs about the feature complexity of the neural representations ^42–44^.

For the EEG-based analysis, we correlated the DNN RDMs with EEG RDMs across all time points obtained from temporally resolved EEG decoding analysis. This yielded a time course of correlation values for each DNN layer, participant, and masking condition. For the fMRI-based analysis, we correlated the DNN RDMs with RDMs from two ROIs (i.e., EVC and LOC), yielding a correlation value per ROI for each DNN layer, participant, and masking condition.

#### Step 2b: Commonality analysis - shared variance among EEG, fMRI and DNN RDMs

To investigate the temporal dynamics of specific visual features emerging in brain regions, we extended standard RSA to commonality analysis ^68,69^ (Fig. 4A). Specifically, we computed the coefficients of shared variance separately among EEG RDMs at each time point, fMRI RDMs in each ROI, and DNN RDMs for each layer. This resulted in a time course of shared variance (R^2^) for each DNN layer, ROI, participant, and masking condition.

To investigate where in the brain the specific visual features originate and how each of the four spectro-temporally identified components carries them, we conducted commonality analysis once more (Fig. 6A). Here, we calculated coefficients of shared variance among frequency-based EEG RDMs corresponding to each spectro-temporally identified component, fMRI RDMs within each ROI, and DNN RDMs across each layer. This analysis resulted in a coefficient of shared variance (R^2^) for each DNN layer, ROI, power- or phase-based component, participant, and masking condition.

#### Noise ceilings

We calculated an upper and lower bound for the noise ceiling ^61^, that is the maximal correlation in the RSA analyses that might be achieved given the noisiness of the data. This was done for the EEG data and fMRI data (i.e., ROIs) separately. To estimate the lower bound, we correlated each participant’s RDM with the average RDM of all other participants. To estimate the upper bound, we correlated each participant’s RDM with the average RDM of all participants. We averaged the results, thus obtaining estimates of the lower and upper noise ceilings for each EEG time point or time-point and frequency combination, as well as for all fMRI ROIs.

### Statistical analyses

We used sign permutation tests ^171^ that do not make assumptions about the data distribution. We compared the statistic of interest (i.e., mean decoding accuracy, correlation coefficients in RSA, coefficients of shared variance in commonality analysis, or differences therein between the masking conditions) against the null hypothesis that the statistic of interest was equal to chance (i.e., 50 % decoding accuracy for pair-wise decoding, a correlation of 0, a coefficient of shared variance of 0, or a difference of 0). To obtain a null distribution, we multiplied participant-specific data randomly by either +1 or -1 and computed the statistic of interest for 10,000 permutations. Based on these null distributions, we obtained p-values by comparing the original statistic to the null distribution. We conducted one-tailed (i.e., the right-tailed) tests for all statistics of interest except for differences, for which we used two-tailed tests.

To correct for multiple comparisons with a small number of unrelated comparisons, we used FDR correction at a p < 0.05 ^172^. In cases involving a large number of comparisons in contiguous and correlated results (i.e., time points, frequencies, or voxels), we used cluster-based inference ^173^. For the cluster-size-based inference, we calculated the statistic of interest both for the empirical results and for each permutation sample under the null hypothesis. This resulted in 1- dimensional (e.g., decoding time courses, RSA-based correlation time courses, time courses of shared variance), 2-dimensional (e.g., decoding time-time matrices, decoding time-frequency matrices, RSA-based correlation matrices), or 3-dimensional (i.e., fMRI volumetric decoding results) p-value maps. We defined clusters based on temporal or spatial contiguity with a p < 0.005 (i.e., cluster-definition threshold) for most analyses, except for time-frequency decoding matrices, which used a threshold of p < 0.05. We determined the maximum cluster size for each permutation sample, yielding a distribution of the maximum cluster size statistic. We set the cluster threshold at p < 0.05.

We calculated 95% confidence intervals for the peak latencies in the resulted time courses (e.g., decoding time courses, RSA-based correlation time courses, time courses of shared variance). For this, we randomly sampled participants with replacements 1,000 times. For each bootstrap sample, we determined the peak latency. This yielded a distribution of peak latencies for which we report the 95 % confidence intervals.

## Acknowledgments

We are grateful to Marleen Haupt and Agnessa Karapetian for their valuable comments on the manuscript, to Lixiang Chen for his assistance with data collection, and to all the participants for their time and contributions to this study. We thank the HPC service of FUB-IT, Freie Universitaet Berlin, for computing time on Curta (http://dx.doi.org/10.17169/refubium-26754). R.M.C. is supported by the German Research Foundation (DFG; CI241/1-1, CI241/3-1, CI241/7- 1) and by a European Research Council (ERC) starting grant (ERC-2018-STG 803370). D.K. is supported by the German Research Foundation (DFG; SFB/TRR135, project number 222641018; KA4683/5-1, project number 518483074), “The Adaptive Mind”, funded by the Excellence Program of the Hessian Ministry of Higher Education, Science, Research and Art, and an ERC Starting Grant (PEP, ERC-2022-STG 101076057).

## Author contributions

Conceptualization: S.X., J.S., D.K. and R.M.C.; Methodology: S.X., J.S., D.K. and R.M.C.; Software: S.X.; Formal analysis: S.X. and J.S.; Investigation: S.X. and B.Y.; Resources: S.X. and R.M.C.; Data curation: S.X.; Writing - original draft preparation: S.X. and R.M.C.; Writing - review and editing: S.X., J.S., D.K. and R.M.C.; Visualization: S.X.; Supervision: R.M.C. and D.K.; Project administrations: R.M.C. and D.K.; Funding acquisition: R.M.C. and D.K.

## Declaration of interests

The authors declare no competing interests.

## Supplementary information

Document S1. Supplementary Discussion 1, Supplementary Figures 1-7 and Supplementary Tables 1-7.

## Notes

### Competing Interest Statement

The authors have declared no competing interest.

