## Supplementary Information for "The representational nature of spatio-temporal recurrent processing in visual object recognition"

### Supplementary Discussions

#### Supplementary Discussion 1 The role of spectro-temporal components in orchestrating visual object processing

Our results show that distinct representations are encoded in spectral power and phase at different time points of visual processing.

In terms of spectral power, efficient masking first affected stimulus representations in a theta/alpha cluster that roughly spans the typical range of time-locked broadband responses (until 300ms post-stimulus, Fig. 5D). This suggests a stimulus-specific modulation of time-locked responses through fast recurrent mechanisms, consistent with the modulation of low-complexity features in EVC that was linked to this cluster (Fig. 6D). This matches a previous report of this spectral signature during object processing using MEG ^1^. Second, we found a later power cluster in the alpha/beta range (emerging after 400ms, Fig. 5D), where representations were different for the early and late mask conditions. This effect is consistent with later recurrent processing extracting more high-level object representations, as indexed by our DNN analysis, which linked representations in this alpha/beta cluster to activity in LOC and to DNN features of higher complexity (Fig. 6G). This delayed involvement of low-frequency rhythms in the formation of more complex visual information is consistent with the idea that representations of higher-level, semantic attributes are related to alpha/theta rhythms that emerge after the initial feedforward response ^2^.

In terms of spectral phase, we found a temporally widespread cluster in the theta frequency range, in which representations were disrupted by efficient masking (Fig. 5G). What is the origin of this effect? We speculate that representations of different visual objects may yield different characteristic phase profiles that result from recurrent interactions between stimulus-selective neural populations at different levels of the hierarchy ^1,3,4^. Efficient masking interferes with these recurrent loops and may thereby reduce the stimulus-information retrievable from these phase components. Interestingly, we also found a more transient phase cluster (from around 100 –400ms post-stimulus, Fig. 5G) in the alpha/beta range that extended into gamma frequencies, in which stimulus information was less prominent when the stimulus was effectively masked. This may again index fast recurrent processes that are more strongly phase-locked to stimulus on- and offset than later recurrent processes. We speculate that this cluster contains a blend of feedforward processing (in gamma frequency range) and feedback processing (in the alpha/beta range), indexing an exchange of information in feedforward-feedback loops that unfold in stimulus-specific phase dynamics ^1^. These early recurrent loops may stabilize the representations of low- and mid-complexity features in early visual cortex (Fig. 6D).

### Supplementary Figures

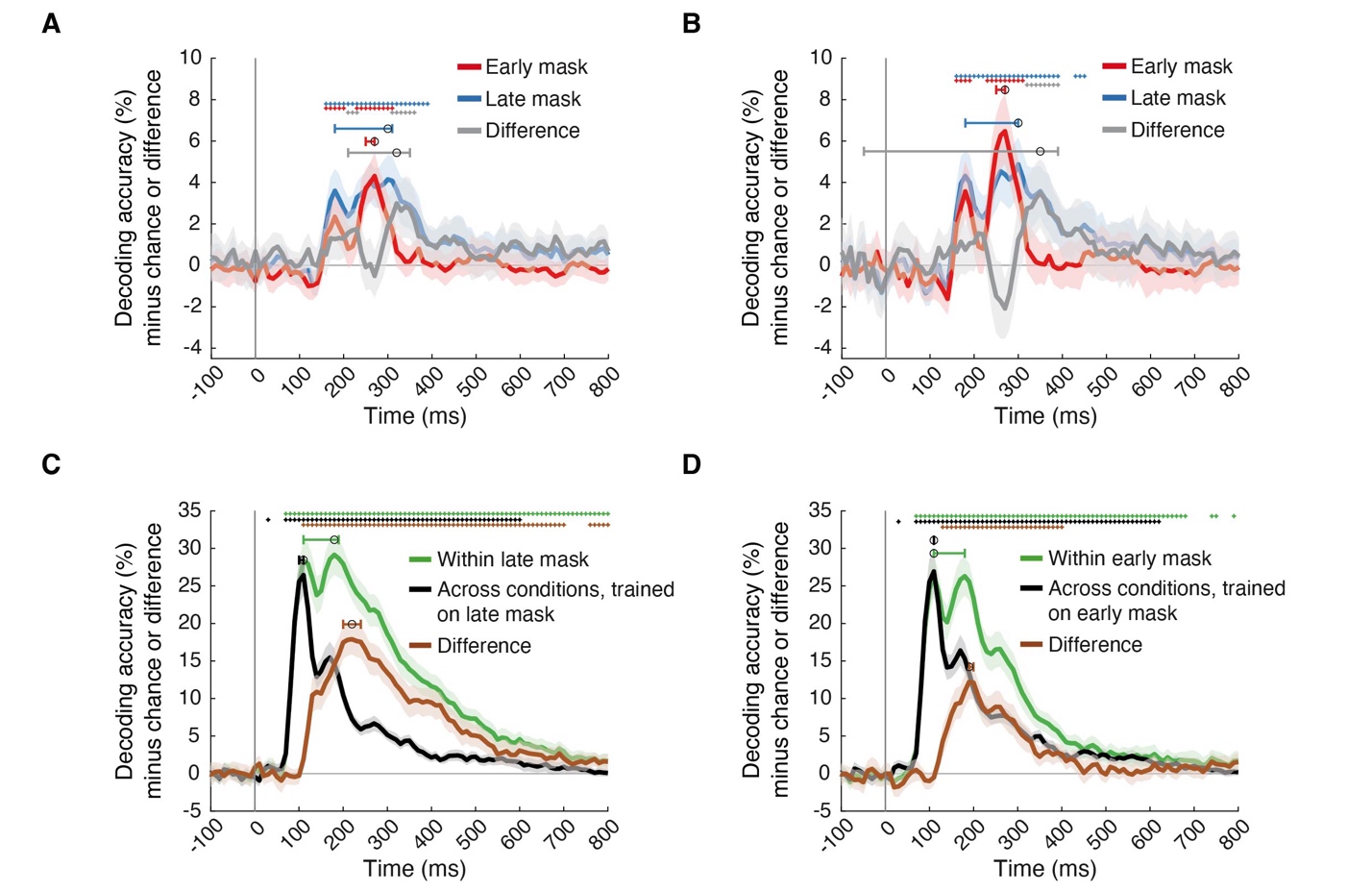

#### Supplementary Figure 1 Temporal dynamics of visual object representations for the two masking conditions.

**(A, B)** Temporal dynamics of object representations across categorical boundaries of naturalness **(A)** and animacy **(B)**. **(C, D)** Pairwise object identity decoding results within (green) and across masking conditions (black), along with their differences (brown), presented separately for the **(C)** late mask condition and **(D)** the early mask condition. Cross-classification results are sorted by training set. For **(A-D)**, decoding chance level was 50%; significant above-chance level decoding is denoted by colored asterisks at the corresponding time points (N = 31, p < 0.05, right-tailed permutation tests, cluster definition threshold p < 0.005, cluster-threshold p < 0.05, 10,000 permutations); vertical gray line at 0ms indicates stimulus onset; shaded margins of time courses indicate 95% confidence intervals of the decoding performance determined by bootstrapping (1,000 iterations); horizontal error bars indicate 95% confidence intervals for peak latencies.

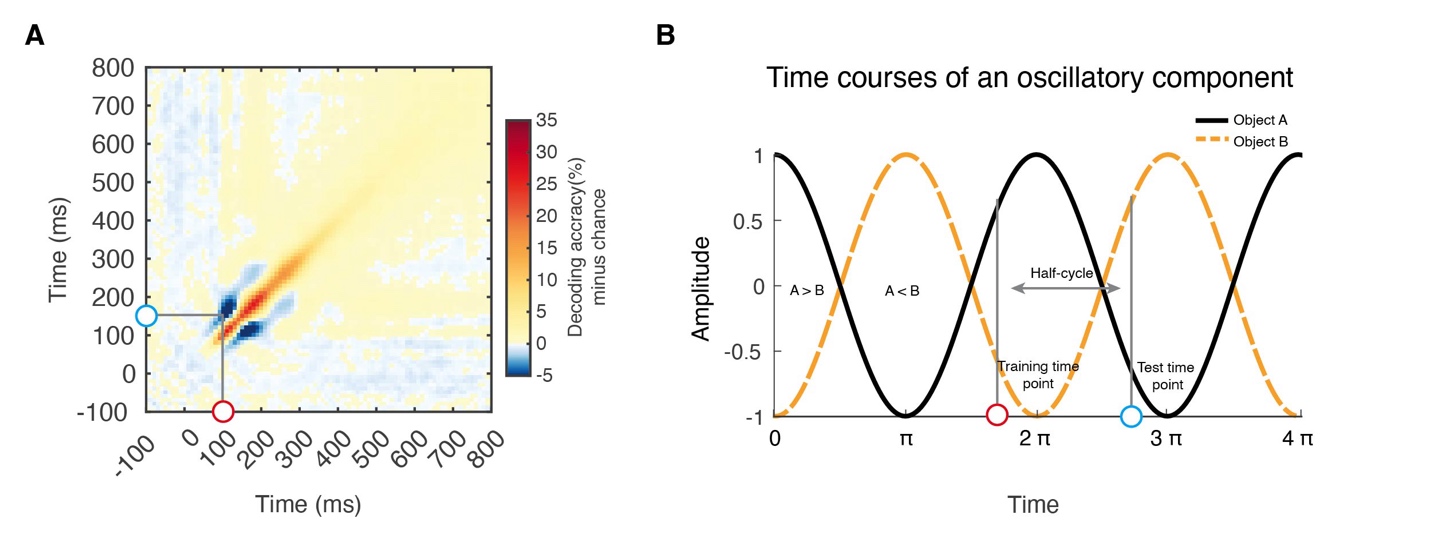

#### Supplementary Figure 2 Potential neural source of negative off-diagonal decoding accuracies in temporal generalization analysis.

**(A)** Results of time-generalized object identity decoding in the early mask condition, as shown in Fig. 1E. Blue and red dots indicate an example pair of corresponding time points for negative off-diagonal decoding accuracies. **(B)** Illustration of a potential oscillatory mechanism underlying negative off-diagonal decoding accuracies. We assume two identical, time-locked responses that differ in phase (here black solid curve and orange dashed curve for object A and B). The time-locked oscillatory responses reverse amplitude with a half-cycle length corresponding to the time difference as determined in **(A)** (i.e., between the corresponding off-diagonal pair of time points, indicated here as in **A** as blue and red circles). This reversal in amplitude results in a reversal in the relationship between the signals for objects A and B (here: at 0π signal A > signal B, at π signal B > signal A). Thus, a classifier trained at time point 0π to distinguish objects A and B based on signal amplitude may predict object identity at time point π with the opposite classification as for time point 0π, resulting in significant below-chance classification accuracy.

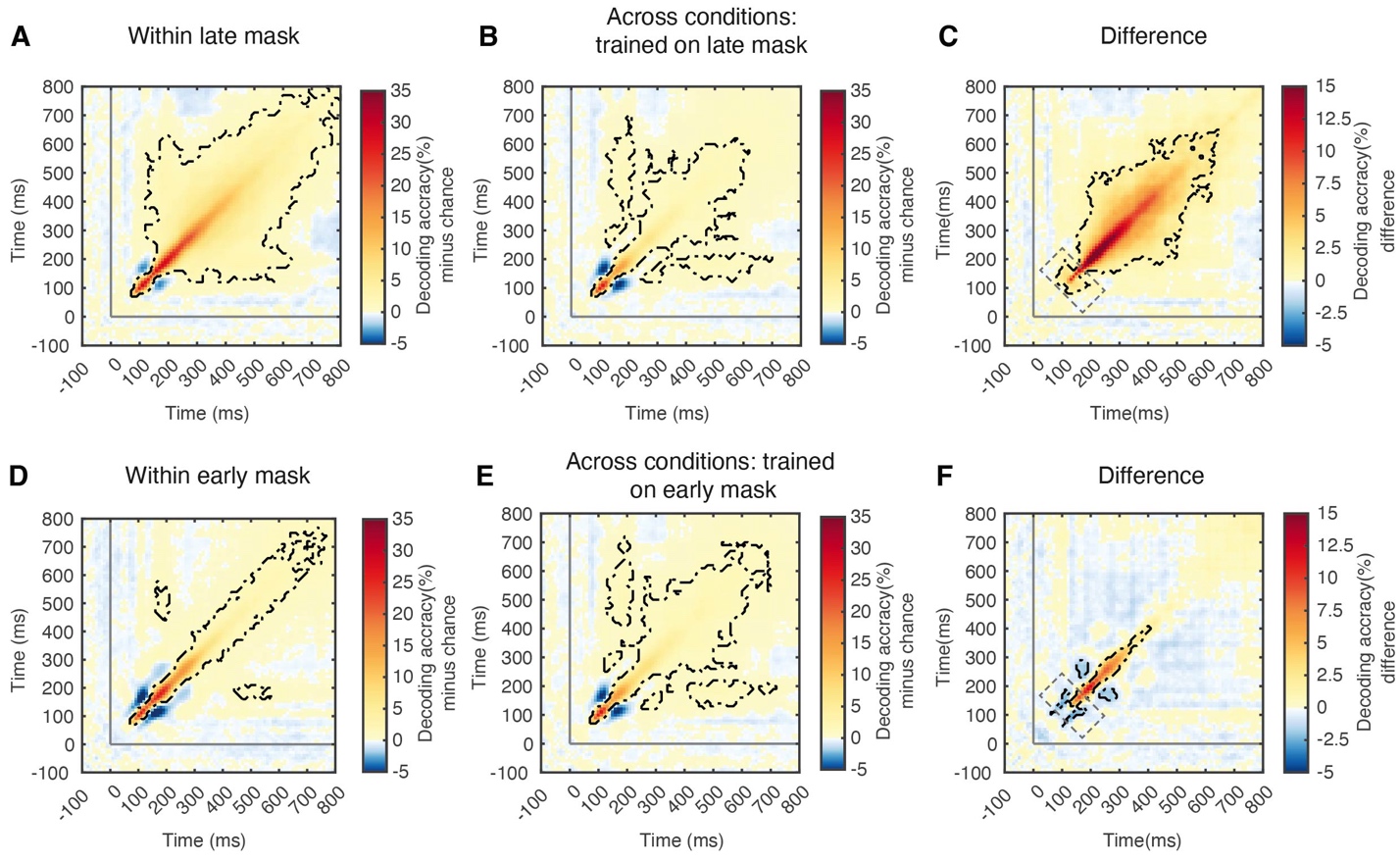

#### Supplementary Figure 3 Results of temporal generalization analysis decoding object identity within- and across-conditions.

**(A)** Results of time-generalized object identity decoding within the late mask condition (same as Fig. 1F). **(B)** Cross-decoding object identity using a classifier trained on the late mask condition. **(C)** The differences between **(A)** and **(B)**. The difference plot reveals positive decoding results in the off-diagonal areas (as shown in the square rectangle in Fig. 1G). This occurs due to higher decoding accuracies in the within-condition decoding (late mask) than the across-conditions decoding (trained on late mask). This confirms the main results pattern: in the within late mask condition decoding **(A)**, the negative off-diagonal decoding results are veiled by recurrent processes. In the across-conditions decoding **(B)**, results are intermediate between the late mask and early mask results. Subtracting the former from the latter results in positive off diagonal decoding accuracies.
**(D)** Results of temporal generalization analysis decoding object identity within the early mask condition (same as Fig. 1E). **(E)** Cross-decoding object identity using a classifier trained on the early mask condition. **(F**) The difference between **(D)** and **(E)**. The difference plot reveals an opposite pattern to the main analysis result (Fig. 1G and **(C)**), with negative decoding results in the off-diagonal areas (as shown in the square rectangle as in Fig. 1G). This occurs due to lower decoding accuracies in the within-condition decoding (early mask) than across-conditions decoding (trained on early mask). This also confirms the main results pattern: in the within early mask condition decoding **(D)**, negative off-diagonal decoding results are not veiled by recurrent processes. In the across-conditions decoding **(E)**, results are intermediate between the late mask and early mask results. Subtracting the former from the latter results in negative off diagonal decoding accuracies.
For **(A-F)**, chance level is 50%. Time-point combinations with significantly above-chance level decoding are outlined in black dash lines (N = 31, right-tailed permutation tests, cluster definition threshold p < 0.005, cluster-threshold p < 0.05, 10,000 permutations); vertical and horizontal gray lines indicate stimulus onsets.

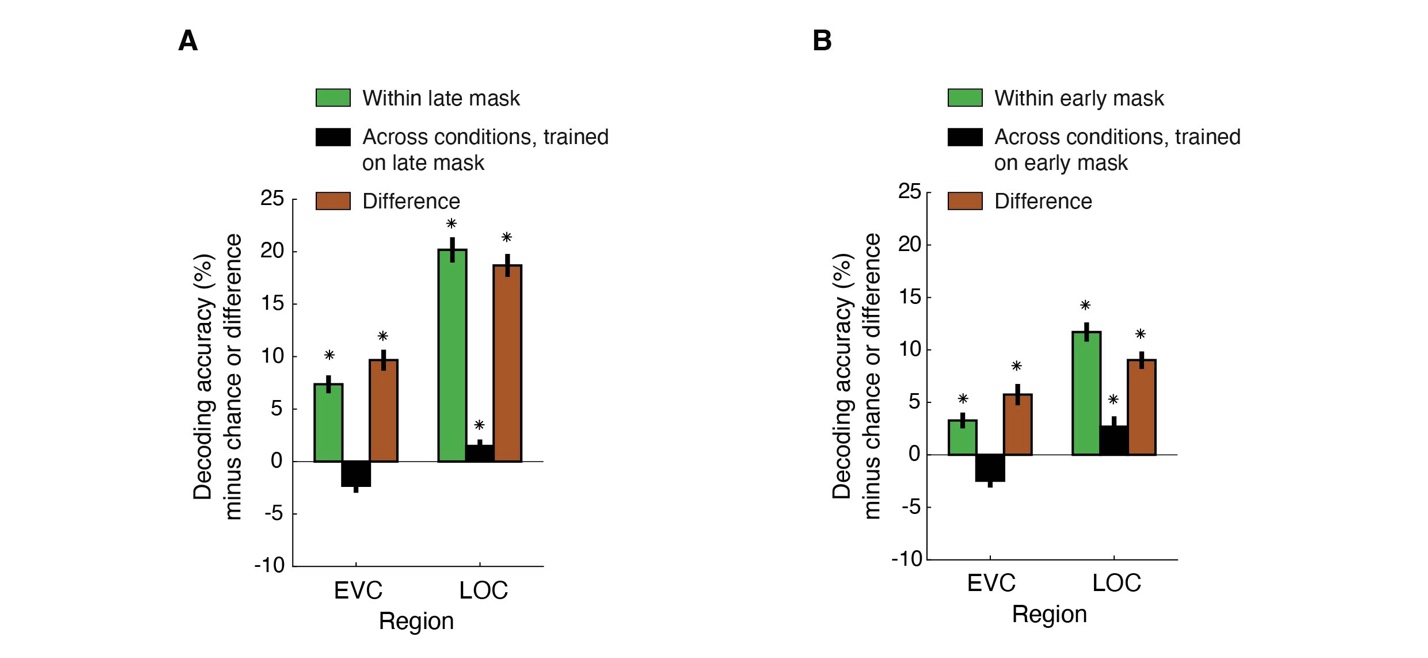

#### Supplementary Figure 4 Results of visual object decoding in fMRI within and across masking conditions.

**(A)** Results of object identity decoding in the late mask condition, across conditions decoding while training on the late mask condition, and the difference between them. **(B)** Results of object identity decoding in the early mask conditions, across conditions decoding while training on the early mask condition, and the difference between them. **(A)** and **(B)** show a qualitatively equivalent results pattern emerges as in Fig. 2C. For **(A, B)**, chance level is 50%; significant above-chance level decoding is denoted by black asterisks above the bars (N = 27, p < 0.05, right-tailed permutation tests, FDR-corrected); error bars indicate standard errors of the mean.

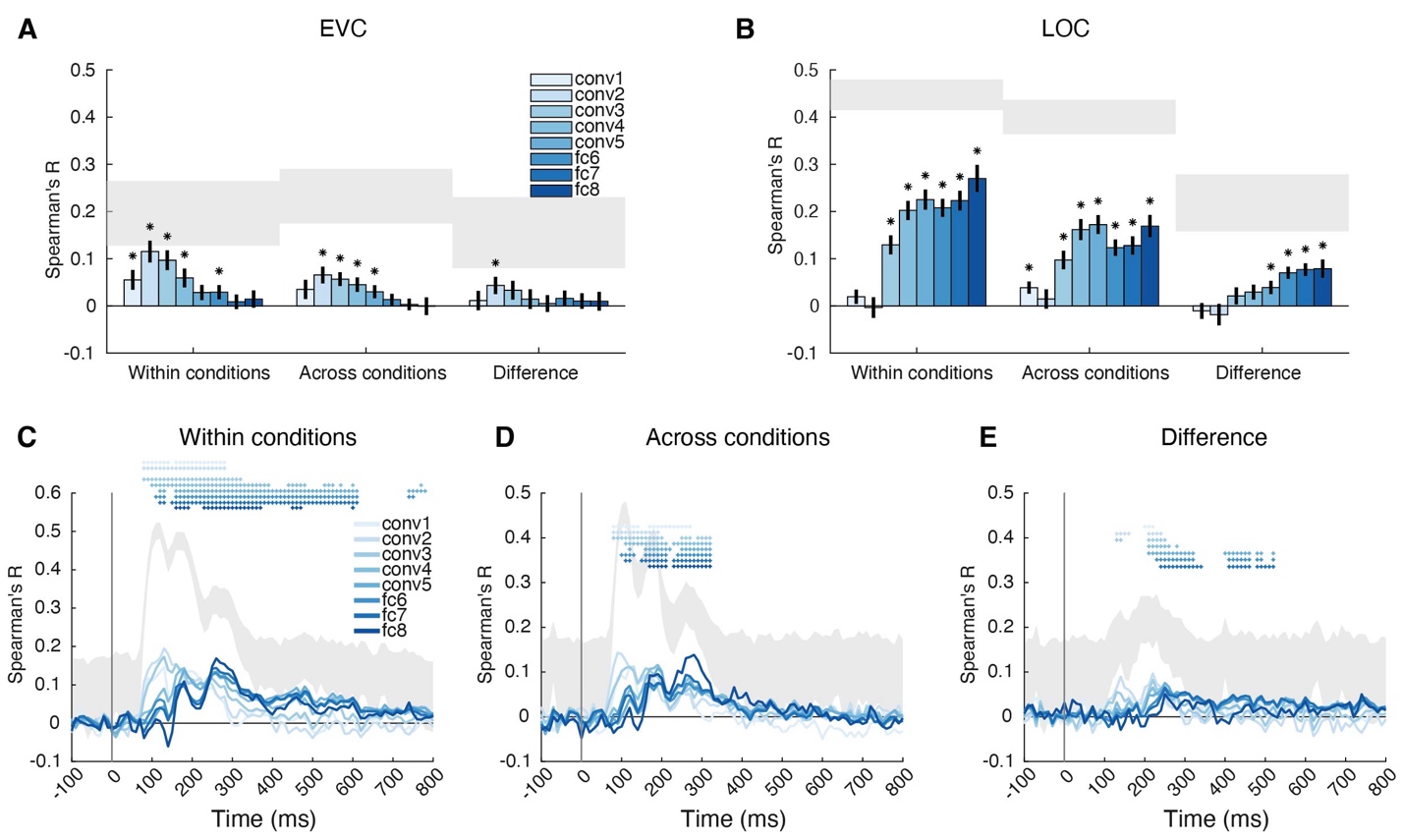

#### Supplementary Figure 5 Visual features encoded in neural object representations revealed by within- and across-conditions decoding.

The analysis rationale is consistent with that used for Fig. 1D and Fig. 2C. We compared the within-conditions decoding results (averaged across within-early-mask and within-late-mask decoding) to the across-conditions decoding results (averaged across both training and testing directions for cross-decoding). This comparison further determines the direct impact of recurrent activity on visual object representations.

**(A, B)** Result of RSA linking object representations in **(A)** EVC and **(B)** LOC to a DNN model trained on object categorization (i.e., AlexNet) as revealed by within condition decoding, across-conditions decoding, and the difference between them. We observe an equivalent result pattern to the main analysis reported in Fig. 3B and 3C. Significant correlations are marked by black asterisks above bars (N = 27, p<0.05, right-tailed permutation tests, FDR corrected); error bars depict standard errors of the mean; shaded gray areas indicate the noise ceiling.
**(C-E)** RSA results linking the DNN to EEG for the **(C)** within condition decoding analysis, **(D)** the across condition analysis, and **(E)** the difference. We observe an equivalent result pattern to the main analysis reported in Fig. 3D-F (for statistical details, see Supplementary Table 4). Significant correlations at time points are denoted by asterisks colored by layer (N = 31, right-tailed permutation tests, cluster definition threshold p < 0.005, cluster-threshold p < 0.05, 10,000 permutations); horizontal error bars indicate 95% confidence intervals for peak latencies, shaded gray areas represented the noise ceiling.

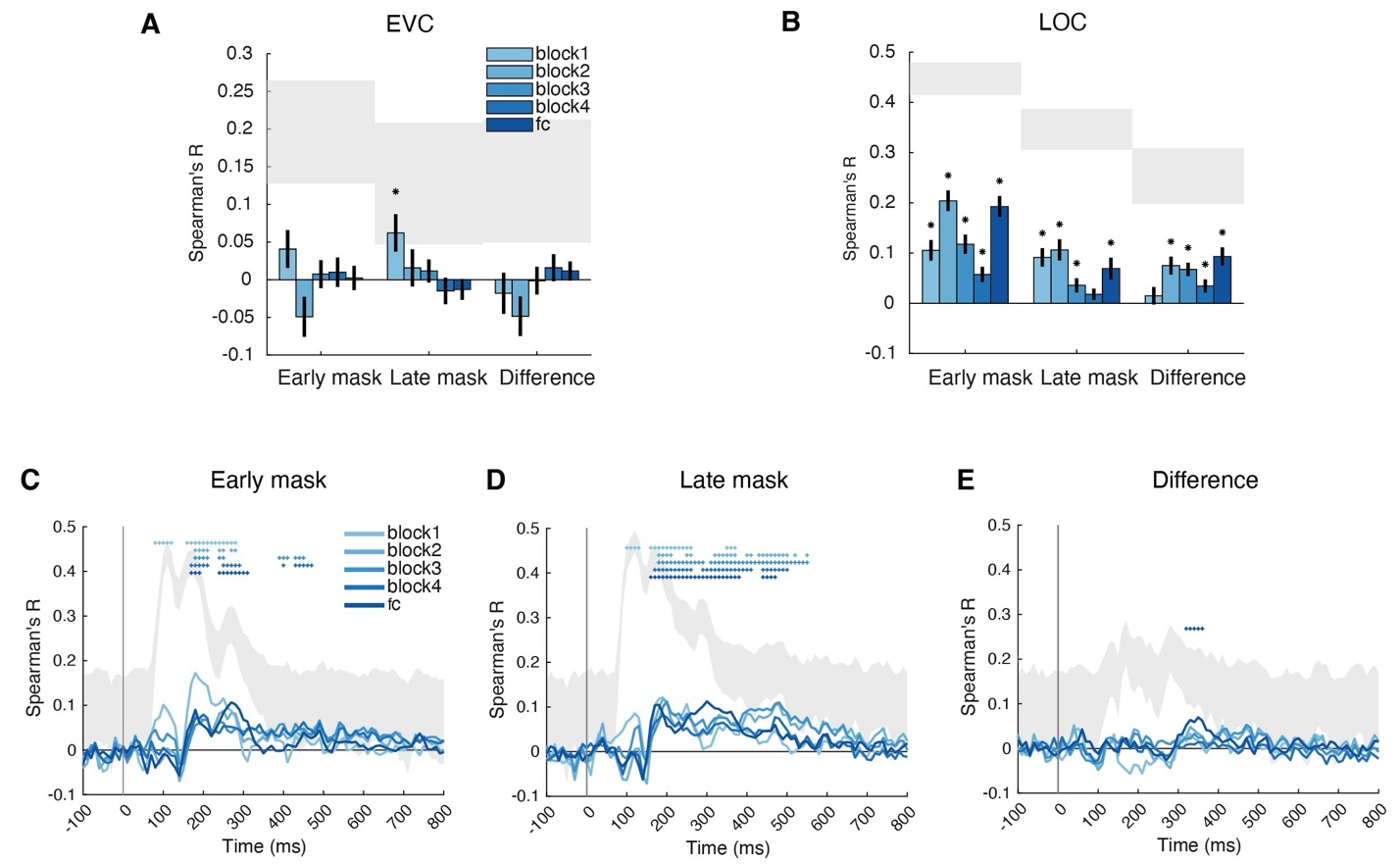

#### Supplementary Figure 6 The representational format of visual representations resolved in space or time as assessed with ResNet50.

We obtained RDMs from the layers of a DNN model (here ResNet50, we used the last layer of each of the four residual blocks and the final classification layer), each ROI in fMRI, and each time point in EEG. We then calculated the correlation coefficients between the DNN layer RDMs and the EEG or fMRI RDMs. **(A, B)** RSA results linking **(A)** EVC and **(B)** LOC to layers of ResNet50. For **(A, B)**, significant correlations are marked by black asterisks above bars (N = 27, p<0.05, right-tailed permutation tests, FDR corrected); error bars depict standard errors of the mean; shaded gray areas indicate the noise ceiling. **(C-E)** RSA results linking layers of ResNet50 to EEG in the **(C)** early mask condition, **(D)** late mask condition, and **(E)** the difference between them. For **(D-F)**, significant correlations at time points are denoted by asterisks colored by layer (N = 31, right-tailed permutation tests, cluster definition threshold p < 0.005, cluster-threshold p < 0.05, 10,000 permutations); horizontal error bars indicate 95% confidence intervals for peak latencies, shaded gray areas represented the noise ceiling.

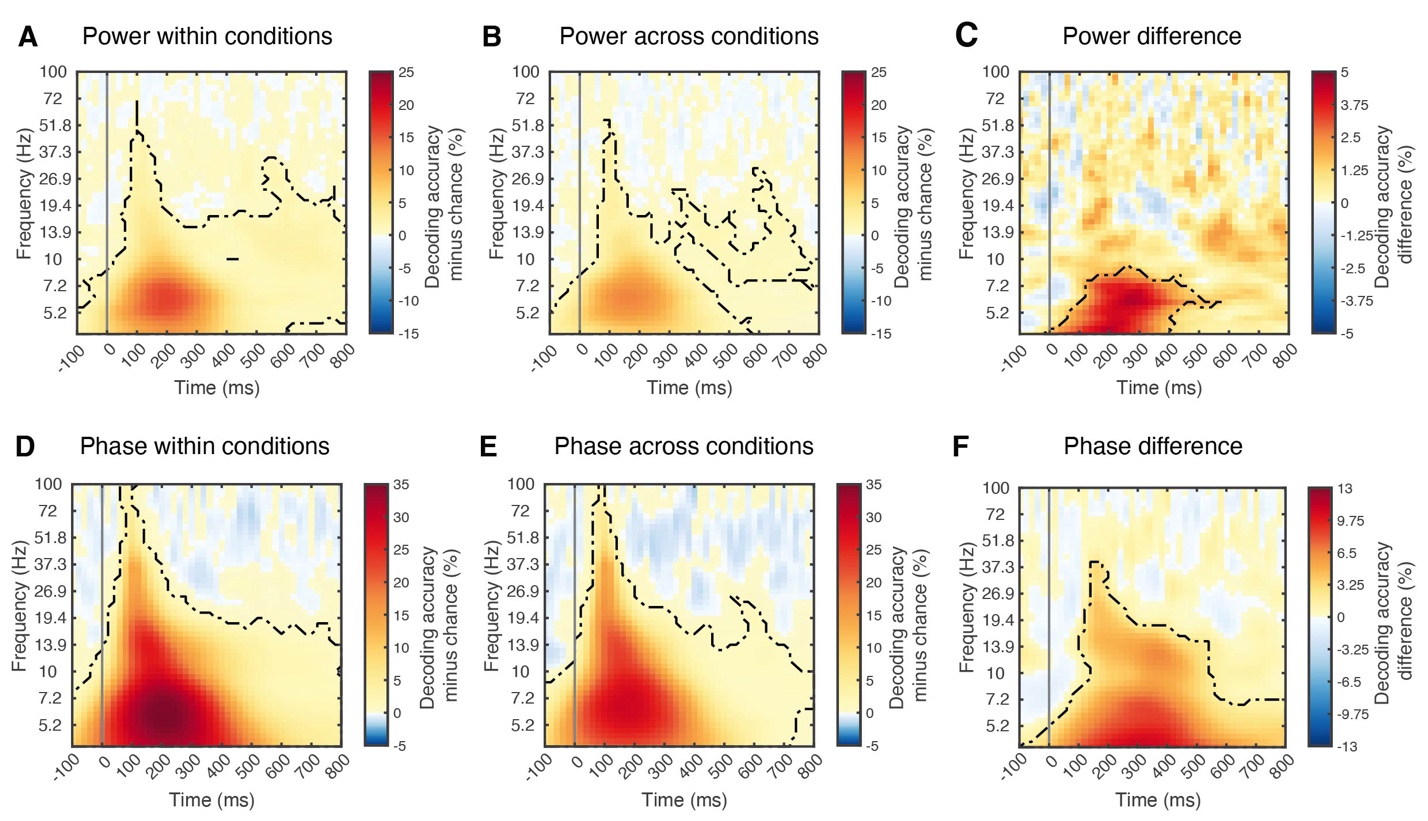

#### Supplementary Figure 7 Spectral characteristics of visual representations as revealed by within and across masking conditions decoding.

The analysis rationale is consistent with that used for Fig. 1D and Fig. 2C. We compared the within-condition decoding results (averaged across within-early-mask and within-late-mask decoding) to the across-condition decoding results (averaged across both training and testing directions for cross-decoding). This comparison helped determine the direct impact of recurrent activity on visual object representations. **(A-F)** Results of time- and frequency-resolved object identity decoding within conditions, across conditions, and their differences. The decoding analyses were based on frequency power values in **(A-C)** and phase values in **(D-F)**. For **(A-F)**, chance level was 50%; time-frequency combinations with significant above-chance decoding are outlined by black dash lines (N = 31, right-tailed permutation tests, cluster definition p < 0.05, significance p < 0.05, 10,000 permutations); the vertical gray line indicates stimulus onset.

### Supplementary Tables

#### Supplementary Table 1 Statistical details for object identity decoding using EEG signals.

| **Type of decoding** | **Peak value*** | **Peak latency (95% CI) #** | **Significant time points+** |
| --- | --- | --- | --- |
| Within-condition (early mask) | 26.42% | 110ms (110, 180) | [70:680, 740:750, 790:800] |
| Within-condition (late mask) | 29.14% | 180ms (110, 190) | [70:800] |
| Difference (within late minus within early mask) | 7.89% | 230ms (220, 420) | [110:560] |
| Across-conditions (averaged across both training and testing directions) | 26.69% | 110ms (100, 110) | [30, 70:600, 620] |
| Difference (within-conditions minus across-conditions decoding) | 14.78% | 200ms (200, 210) | [120:540, 570, 590:650, 680:690, 730:800] |

* Decoding accuracy (%) minus chance level (50%)

\# The unit of time was milliseconds and the 95% confidence intervals added in parentheses were calculated by bootstrapping participants (n = 1,000)

+ Right-tailed cluster-based permutation tests, cluster definition p < 0.005, significance p < 0.05

#### Supplementary Table 2 Statistical details for object naturalness and animacy decoding using EEG signals.

| **Type of decoding** | **Peak value*** | **Peak latency (95% CI) #** | **Significant time points+** |
| --- | --- | --- | --- |
| 1) **Naturalness** | | | |
| Within-condition (early mask) | 4.31% | 270ms (250, 270) | [160:200, 230:310] |
| Within-condition (late mask) | 4.15% | 300ms (180, 310) | [160:390] |
| Difference (within late minus within early mask) | 2.99% | 320ms (210, 350) | [210:230, 310:360] |
| 2) **Animacy** | | | |
| Within-condition (early mask) | 6.47% | 270ms (250, 270) | [160:190, 230:310] |
| Within-condition (late mask) | 4.87% | 300ms (180, 300) | [160:390, 430:450] |
| Difference (within late minus within early mask) | 3.50% | 350ms (-50, 390) | [320:390] |
| 3) **Trained on late mask condition** | | | |
| Across-condition (trained on late mask) | 26.45% | 110ms (100, 110) | [30, 70:600] |
| Difference (within late mask minus across-condition trained on late mask) | 17.90% | 220ms (200, 240) | [-180, 110:700, 760:800] |
| 4) **Trained on early mask condition** | | | |
| Across-condition (trained on early mask) | 26.93% | 110ms (110, 110) | [30, 70:620] |
| Difference (within early mask decoding minus across-conditions trained on early mask) | 12.21% | 190ms (190, 200) | [130:400] |

* Decoding accuracy (%) minus chance level (50%)

\# The unit of time was milliseconds and the 95% confidence intervals added in parentheses were calculated by bootstrapping participants (n = 1,000)

+ Right-tailed cluster-based permutation tests, cluster definition p < 0.005, significance p < 0.05

#### Supplementary Table 3 Statistical details for the RSA results linking the AlexNet model to EEG decoding RDMs within the early and late mask conditions, and the difference between these conditions.

| **Condition**  **Layer group** | **Early mask** | | | **Late mask** | | | **Difference (late mask minus early mask)** | | |
| --- | --- | --- | --- | --- | --- | --- | --- | --- | --- |
|  | Peak value* | Peak latency (95% CI) # | Significant time points+ | Peak value* | Peak latency (95% CI) # | Significant time points+ | Peak value* | Peak latency (95% CI) # | Significant time points+ |
| 1 | 0.13 | 200ms (100, 210) | [80:130, 160:280] | 0.14 | 130ms (90, 130) | [70:140, 160:270] | n.s. | n.s. | n.s. |
| 2 | 0.15 | 110ms (100, 140) | [80:280] | 0.18 | 130ms (100, 160) | [80:260, 280] | n.s. | n.s. | n.s. |
| 3 | 0.13 | 110ms (110, 220) | [80:320] | 0.17 | 130ms (120, 170) | [80:280] | 0.06 | 160ms (-70, 740) | [160:170] |
| 4 | 0.12 | 260ms (180, 280) | [110:320] | 0.14 | 170ms (160, 190) | [110:400, 430:500, 550:580, 600:610] | 0.08 | 470ms (-100, 610) | [160:170, 470] |
| 5 | 0.12 | 260ms (180, 280) | [160:340] | 0.13 | 180ms (170, 480) | [120:130, 160:610] | 0.07 | 470ms (-100, 480) | [170, 360, 470:480] |
| 6 | 0.12 | 260ms (250, 280) | [160:190, 220:330] | 0.10 | 170ms (170, 480) | [110:130, 150:610] | 0.07 | 350ms (140, 480) | [170, 340:380, 470:480] |
| 7 | 0.13 | 260ms (250, 280) | [160:180, 230:330] | 0.11 | 170ms (170, 480) | [120:130, 150:610] | 0.08 | 470ms (340, 480) | [170, 320:370, 440:480] |
| 8 | 0.16 | 270ms (250, 290) | [170:180, 240:310] | 0.11 | 260ms (180, 360) | [160:190, 230:380, 430:480, 600] | 0.11 | 350ms (-110, 470) | [330:360] |

* Spearman correlation coefficients

\# The unit of latency was milliseconds and the 95% confidence intervals added in parentheses were calculated by bootstrapping participants (n = 1,000)

+ Right-tailed cluster-based permutation tests, cluster definition p < 0.005, significance p < 0.05

#### Supplementary Table 4 Statistical details for the RSA results linking the AlexNet model to EEG decoding RDMs within-, across-conditions, and the difference between them.

| **Condition**  **Layer** | **Within conditions** | | | **Across conditions** | | | **Within - across** | | |
| --- | --- | --- | --- | --- | --- | --- | --- | --- | --- |
|  | Peak value* | Peak latency (95% CI) # | Significant time points+ | Peak value* | Peak latency (95% CI) # | Significant time points+ | Peak value* | Peak latency (95% CI) # | Significant time points+ |
| 1 | 0.15 | 130ms (90, 210) | [80:140, 160:280] | 0.11 | 120ms (90, 190) | [80:140, 170:270] | 0.06 | 210ms (-80, 660) | [200:220] |
| 2 | 0.20 | 130ms (120, 140) | [80:280] | 0.14 | 90ms (90, 130) | [70:190] | 0.10 | 220ms (130, 230) | [130:160, 210:240] |
| 3 | 0.17 | 130ms (120, 160) | [80:320] | 0.11 | 120ms (100, 190) | [80:200. 250:270, 290:320] | 0.08 | 220ms (130, 280) | [130:140, 210:250] |
| 4 | 0.14 | 180ms (160, 280) | [100:390, 440:480, 530:550, 570, 600, 770] | 0.12 | 190ms (170, 270) | [110:210, 230:320] | 0.07 | 220ms (160, 680) | [210:260, 280] |
| 5 | 0.14 | 180ms (180, 280) | [120,130, 160:610, 740:780] | 0.11 | 190ms (170, 270) | [120, 160:220, 250:320] | 0.08 | 250ms (230, 450) | [210:320, 400:460, 480:490, 520] |
| 6 | 0.13 | 260ms (250, 290) | [110:130, 160:610, 740:750] | 0.08 | 200ms (120, 290) | [100:130, 160:210, 240:320] | 0.07 | 250ms (240, 680) | [230:320, 410:460, 480:520] |
| 7 | 0.14 | 260ms (250, 290) | [120:130, 150:610] | 0.09 | 260ms (170, 290) | [110:120, 150:210, 230:320] | 0.06 | 250ms (250, 600) | [240:340, 410:460, 480:520] |
| 8 | 0.17 | 260ms (250, 300) | [160:190, 230:370, 450:470, 600] | 0.14 | 280ms (190, 290) | [170:210, 230:320] | n.s. | n.s. | n.s. |

* Spearman correlation coefficients

\# The 95% confidence intervals added in parentheses were calculated by bootstrapping participants (n = 1,000)

+ Right-tailed cluster-based permutation tests, cluster definition p < 0.005, significance p < 0.05

#### Supplementary Table 5 Statistical details for RSA-based commonality analysis results linking RDMs from EEG, fMRI and the AlexNet model.

| **Condition**  **Layer group** | **Early mask** | | | **Late mask** | | | **Difference (late mask minus early mask)** | | |
| --- | --- | --- | --- | --- | --- | --- | --- | --- | --- |
|  | Peak value* | Peak latency (95% CI) # | Significant time points+ | Peak value* | Peak latency (95% CI) # | Significant time points+ | Peak value* | Peak latency (95% CI) # | Significant time points+ |
| 1) **EVC** | | | | | | | | | |
| 1 | 0.005 | 100ms (-30, 510) | [70:130, 190:210, 490:510, 610, 750:760] | 0.007 | 130ms (90, 130) | [90:150] | n.s. | n.s. | n.s. |
| 2 | 0.008 | 120ms (100, 760) | [80:130, 190:200, 410, 450, 500, 590:610, 650:660, 680, 700:710,740:770] | 0.018 | 130ms (90, 130) | [80:130, 200:240, 620:640] | 0.011 | 130ms (80, 160) | 130 |
| 3 | 0.006 | 110ms (100, 580) | [80:130, 190:200, 410:450, 500:530, 580:600, 640:660, 680:760] | 0.014 | 130ms (100, 130) | [70:140, 200:240, 620:630] | 0.010 | 130ms (70, 170) | [110:130] |
| 4 | 0.003 | 110ms (100, 550) | [80, 100:130, 420:430, 450, 510:520, 580, 650:660, 750:760] | 0.005 | 130ms (90:420) | [90, 110:130, 210:220] | 0.003 | 130ms (10, 430) | 130 |
| 5 | 0.001 | 200ms (-30, 520) | [100, 130, 430] | 0.002 | 120ms (110, 180) | [110:130] | n.s. | n.s. | n.s. |
| 6 | n.s. | n.s. | n.s. | 0.001 | 120ms (110, 180) | [110:130] | n.s. | n.s. | n.s. |
| 7 | n.s. | n.s. | n.s. | n.s. | n.s. | n.s. | n.s. | n.s. | n.s. |
| 8 | n.s. | n.s. | n.s. | n.s. | n.s. | n.s. | n.s. | n.s. | n.s. |
| 2) **LOC** | | | | | | | | | |
| 1 | 0.002 | 180ms (180, 190) | [170:190] | n.s. | n.s. | n.s. | n.s. | n.s. | n.s. |
| 2 | 0.002 | 180ms (160, 180) | [160:190] | n.s. | n.s. | n.s. | n.s. | n.s. | n.s. |
| 3 | 0.007 | 180ms (170, 250) | [160:210, 220, 240:280] | 0.011 | 180ms (170, 190) | [160:210] | n.s. | n.s. | n.s. |
| 4 | 0.011 | 180ms (180, 280) | [160:220, 240:280] | 0.019 | 180ms (170. 190) | [160:220, 250, 340:350, 440] | 0.009 | 170ms (160, 480) | [170, 190] |
| 5 | 0.008 | 180ms (180, 280) | [170:190, 210, 240:280] | 0.018 | 180ms (180, 190) | [160:260, 320:370, 400, 430:450] | 0.011 | 190ms (170, 480) | [170:190, 330:340, 360:370, 440:450] |
| 6 | 0.005 | 270ms (180, 280) | [170:180, 250:280] | 0.011 | 170ms (170, 480) | [160:190, 220, 240:300, 320:360, 400:410, 430:450] | 0.009 | 340ms (170, 710) | [160:170, 190, 320, 340:360, 440:450] |
| 7 | 0.004 | 250ms (180, 280) | [170:180, 250:280] | 0.011 | 180ms (170, 480) | [160:190, 220, 240:300, 320:360, 400:410, 430:450] | 0.009 | 170ms (160, 710 | [160:190, 320:360, 440:450] |
| 8 | 0.008 | 270ms (180, 280) | [170:180, 240:280] | 0.017 | 340ms (170, 710) | [170:180, 240:250, 270:280, 300:380, 400, 440:450] | 0.015 | 340ms (170, 710) | [320:380] |

* Coefficients of shared variance

\# The 95% confidence intervals added in parentheses were calculated by bootstrapping participants (n = 1,000)

+ Right-tailed cluster-based permutation tests, cluster definition p < 0.05, significance p < 0.05

#### Supplementary Table 6 Statistical details for object identity decoding using spectro-temporally resolved EEG signals.

| **Type of decoding** | **Peak value *** | **Peak latency (95% CI) #** | **Peak frequency (95% CI) #** | **Significant time points +** | **Significant frequency ranges +** |
| --- | --- | --- | --- | --- | --- |
| 1) **Frequency power values** | | | | | |
| Within-condition (early mask) | 15.00% | 200ms (180, 220) | 6.77hz (5.56, 7.22) | [-140:800] | [4.00:63.14] |
| Within-condition (late mask) | 17.68% | 200ms (180, 200) | 6.34hz (5.20, 6.77) | [-120:800] | [4.00:39.86] |
| Difference (late mask minus early mask) | 5.21% | 540ms (120, 580) | 10.72hz (4, 26.88) | [0:320, 420:800] | [4.00:32.73] |
| 2) **Frequency phase values** | | | | | |
| Within-condition (early mask) | 35.48% | 200ms (200, 220) | 5.93hz (5.56, 6.77) | [-300:800] | [4.00:100.00] |
| Within-condition (late mask) | 36.06% | 200ms (200, 220) | 5.56hz (4.87, 6.77) | [-300:800] | [4.00:100.00] |
| Difference (late mask minus early mask) | 8.29% | 180ms (160, 220) | 20.67hz (16.97, 23.57) | [-20:800] | [4.00:72.00] |

* Decoding accuracy (%) minus chance level (50%)

\# The 95% confidence intervals added in parentheses were calculated by bootstrapping participants (n = 1,000)

+ Right-tailed cluster-based permutation tests, cluster definition p < 0.05, significance p < 0.05

#### Supplementary Table 7 Behavioral performance during EEG and fMRI experiments.

| **Measure**  **Condition** | **EEG experiment** | | | **fMRI experiment** | | |
| --- | --- | --- | --- | --- | --- | --- |
|  | Mean (S.D.) | Difference (95% CI) | P-value* | Mean (S.D.) | Difference (95% CI) | P-value* |
|  | **Correctness minus chance (%) #** | | | **d-prime** | | |
| Overall | 33.27 (6.91) | | | 3.27 (2.26) | | |
| Early mask | 29.03 (8.67 | 8.48 (6.51, 10.54) | <0.001 | 2.01 (1.01) | 2.51 (1.82, 3.32) | <0.001 |
| Late mask | 37.51 (6.11)) |  |  | 4.53 (2.41) |  |  |
|  | **Reaction time (ms)** | | | **Reaction time (ms)+** | | |
| Overall | 504.56 (92.86) | | | 631.62 (154.06) | | |
| Early mask | 513.83 (110.09) | 18.23 (8.10, 30.11) | 0.002 | 652.48 (123.64) | 41.72 (-1.69, 85.37) | 0.029 |
| Late mask | 495.60 (86.73) |  |  | 610.76 (179.74) |  |  |

\# Correctness was the corrected response accuracy (i.e., raw response accuracy (%) minus chance accuracy 50%) for the two alternatives-forced choices task following the presentation of the early mask or the late mask conditions in the EEG experiment

+ Participant number was 24 with 3 missing data

* P-values denoted significance level of difference by paired t-tests (N=31 in EEG experiment, N=27 in fMRI experiment) comparing the mean of the measurements taken from the same participant between early mask and late mask conditions
